# Mitosis exit followed by death in interphase prevents the development of polyploid giant cancer cells

**DOI:** 10.1101/2023.08.31.555795

**Authors:** Juan Jesus Vicente, Kainat Khan, Grant Tillinghast, José L. McFaline-Figueroa, Yasemin Sancak, Nephi Stella

## Abstract

Microtubule targeting agents (**MTAs**) are commonly prescribed to treat cancers and predominantly kill cancer cells in mitosis. Significantly, some MTA-treated cancer cells can escape death in mitosis and exit mitosis, and become malignant polyploid giant cancer cells (**PGCC**). Considering the low number of malignant cells undergoing mitosis in tumor tissue, killing these cells in interphase may represent a favored antitumor approach. We discovered that ST-401, a mild inhibitor of microtubule assembly, preferentially kills cancer cells in interphase as opposed to mitosis, and avoids the development of PGCC. Single cell RNA sequencing identified mRNA transcripts regulated by ST-401, including mRNAs involved in ribosome and mitochondrial functions. Accordingly, ST-401 induces an integrated stress response and promotes mitochondria fission accompanied by a reduction in energy metabolism. This cell response may underly death in interphase and avoid the development of PGCC.

## Introduction

Most microtubule targeting agents (**MTAs**) bind directly to either tubulin dimers or assembled microtubules (**MTs**), and depending on their mechanism of action, may inhibit either MT assembly or disassembly ^1, 2^. Structural biology studies have identified at least seven ligand-binding sites on tubulin for MTAs, including sites targeted by small molecular inhibitors of MT assembly such as colchicine, vincristine and eribulin ^3, 4^. In the case of agents that bind to the colchicine site, these inhibitors of MT assembly bind directly to either tubulin dimers or assembled microtubules (**MTs**), and depending on their mechanism of action inhibit either MT assembly or disassembly ^1, 2^. Significantly, proper operation of dynamic MTs within the spindle is essential to achieve error-free mitosis. Thus, abnormal changes in MT dynamics induced by MTAs will keep the spindle assembly checkpoint (**SAC**) activated and halt cell cycle progression of metaphase cells (metaphase arrest), resulting in a delay of anaphase onset until all chromosomes have acquired proper attachments to MTs ^5^. Consequently, extended metaphase arrest will ultimately trigger cell death through different mechanisms, including apoptosis, necrosis, and autophagy ^6^. Landmark studies demonstrated that most cancers exhibit slow doubling times and low mitotic indices (i.e., below 5%); and yet MTAs have proven remarkably efficacious antitumor agents ^4, 7^. This premise led the field to hypothesize that MTAs might also kill cancer cells in interphase and few examples have been reported ^8, 9^; yet the molecular mechanism by which cancer cell death in interphase occurs remains unknown.

Cancer cells treated with potent inhibitors of MTA assembly, such as nocodazole (**NOC**), which binds to the colchicine site of tubulin, predominantly die during mitosis because of pronounced disruption of the mitotic spindle machinery; however, cells that escape SAC-induced death, exit mitosis with different levels of chromosome missegregation to produce a plethora of daughter cells with different levels of aneuploidy, among them malignant polyploid giant cancer cells (**PGCCs**) ^10^. Specifically, live microscopy tracking of cell fate experiments revealed that mitotically arrested cells fail to undergo cytokinesis and exit mitosis through mitotic slippage to become PGCCs. Evidence demonstrates that PGCCs carry cancer-stem-cell features and can be implicated in increased malignancy and enhanced metastatic characteristics, including drug resistance ^11–15^. These studies favor the concept that killing cancer cells in interphase may represent a safer and more efficacious antitumor approach than targeting mitosis. However, understanding the mechanism involved in cell death in interphase has been difficult to resolve owing to the inability to specifically isolate this event with selective compounds and to identify the time course of this cellular response.

In previous work, we developed a series of carbazole-based compounds (*ST-compounds*) that bind to the colchicine site of tubulin with high-affinity and reversibly reduce MT assembly rates ^16–19^ (**Figure 1a**). Live fluorescence microscopy imaging to track assembling MTs in cancer cells in interphase demonstrated that nanomolar concentrations of ST-401, our most potent compound, triggered a milder reduction in MT assembly as compared to nanomolar concentrations of NOC ^19^. This milder response occurred even though nanomolar concentrations of ST-401 and NOC are equally effective and exhibit similar IC_50_s at inhibiting MT assembly in biochemical assays ^19^. Another difference was that MTs were still visibly intact in cancer cells in interphase treated with low nanomolar concentrations of ST-401 as compared to NOC treatment ^19^. Together, these results suggested a milder response triggered by ST-401 compared to NOC and raised the possibility that ST-401 might induce a different antitumor response than NOC. Based on this premise, here we evaluated the antitumor activity of ST-401 in the NCI-60 cancer cell line panel and used live-cell microscopy imaging to identify the time course of cell death induced by ST-401 compared to NOC. We then used single cell RNA sequencing to identify the molecular mechanisms that are differentially regulated by ST-401 versus NOC, thus identifying molecular mechanisms that result in the killing of cancer cells in interphase and prevent the formation of PGCCs. We show that ST-401 induces an integrated stress response (**ISR**) and promotes mitochondrial fission, accompanied by a reduction in energy metabolism, cellular responses that may underly death in interphase and avoid the development of PGCCs.

**Figure 1:**
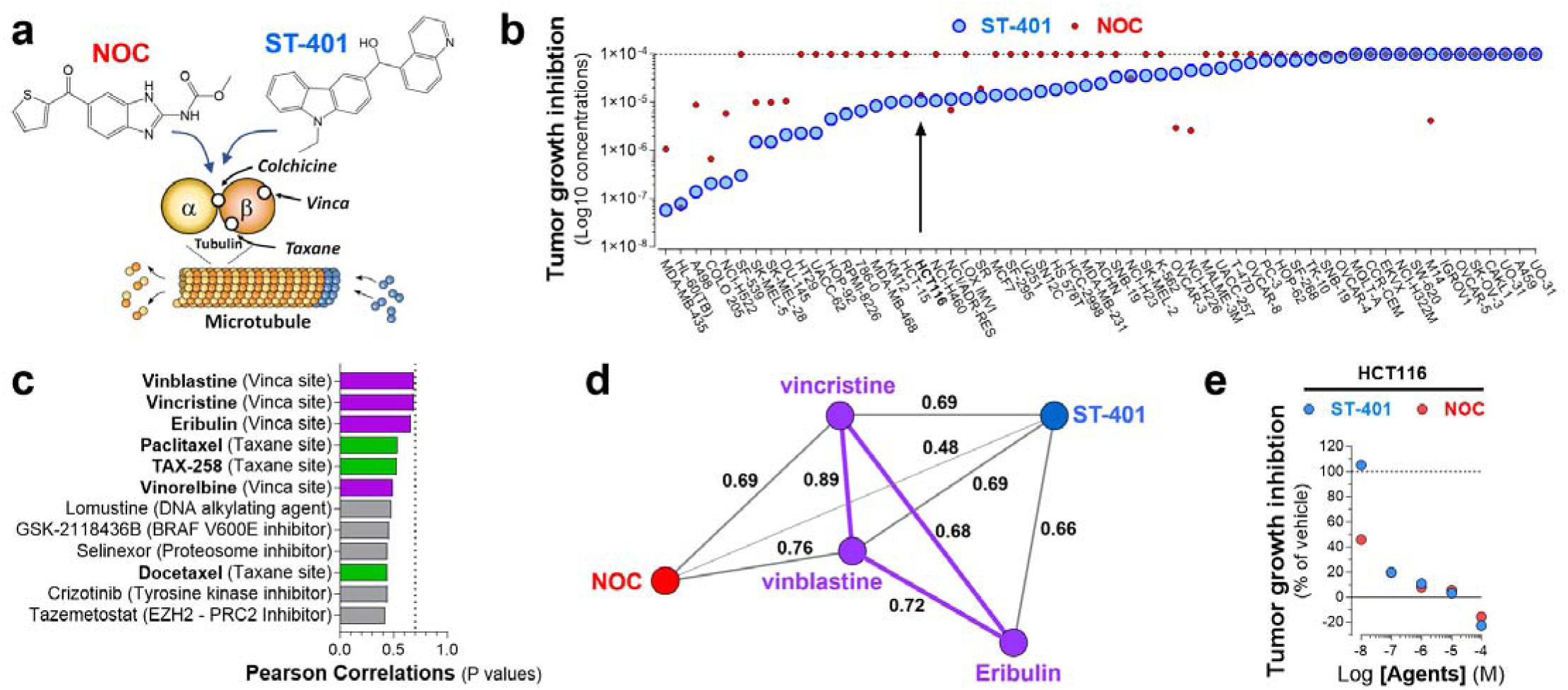
ST-401 and NOC kill cancer cells through a different antitumor mechanism: evidence from the NCI-60 cancer cell line panel. **a)** Diagram illustrating ST-401 and NOC binding to the colchicine site of tubulin distinct from the vinca and taxane sites, all of which differentially influence MT dynamics via their binding to distinct sites on tubulin. **b)** The antitumor activity of ST-401 was tested in the NCI-60 cancer cell line panel and results analyzed using COMPARE software. ST-401 exhibited significant antitumor activity in 46 out of 60 cancer cell lines as measured by total growth inhibition (TGI) of HCT116 cells exhibited an average sensitivity (arrow). **c)** The top 12 compounds with best correlations in antitumor activity (Pearson correlation) with ST-401. MTAs that target the vinca site (purple) or the taxane site (green) are among the top 12 (7 out of 12). Initial considerations of significance using COMPARE software is normally set at >0.7 for Pearson correlation values (dotted line) and was not reached by any of the top 12 compounds. **d)** Vector map representation of the direct Pearson correlations between ST-401 (blue) and its closest MTAs acting on the vinca site (purple) and nocodazole (NOC) (red). **e)** Direct comparison of the antitumor activity of ST-401 and NOC measured in HCT116 cells within the NCI-60 panel suggests a different response at low nM concentrations.

## Results

### Distinct antitumor activities associated with ST-401 and NOC revealed by the NCI-60 cancer cell line panel analysis

We tested the antitumor activity of ST-401 in the NCI-60 cancer cell line panel and found that it significantly inhibited the growth of 47 out of 60 cancer cell lines as measured by total growth inhibition (**TGI**) (Log_10_ concentrations ranging from 58 nM to 860 μM), showing that cancer cell lines exhibit differential sensitivities to ST-401 (**Figure 1b**). The NCI-60 cancer cell lines were differentially killed by NOC (**Figure 1b**), which was significantly less potent than ST-401. We further analyzed ST-401’s antitumor activity using COMPARE to evaluate whether its antitumor activity correlated with any of the antitumor activity induced by >100,000 chemical compounds and natural products tested on this panel thus far, including 169 compounds in the Approved Oncology Agents library ^20, 21^. **Figure 1c** shows that the antitumor activity of ST-401 did not significantly correlate (i.e., Pearson correlation >0.7) with any compound tested on this platform thus far, and that the initial 12 compounds that resulted in weak correlations included MTAs that bind in distinct sites on tubulin, including vincristine, vinblastine and eribulin, as well as several taxanes, and did not include any MTAs that bind in the colchicine site, such as NOC. **Figure 1d** shows a vector representation of the antitumor activity of ST-401 and how it relates to the antitumor activity induced by compounds that exhibit the closest antitumor activity: namely, the antitumor activity of ST-401 is closer to the antitumor activities of vincristine, vinblastine and eribulin, which act through different sites on tubulin, and is remarkably different than the antitumor activity of NOC. The difference between the antitumor activity of ST-401 and NOC is further emphasized by overall direct correlations between vincristine, vinblastine and eribulin that were higher than their correlation with ST-401 or NOC (**Figure 1d**). Together, these results suggest that ST-401 kills cancer cells through a different mechanism than these MTAs and all Approved Oncology Agents. These results also suggest that ST-401 elicits a significantly different antitumor mechanism than NOC though they both bind in the colchicine site on tubulin.

HCT116 cells are a near-diploid colorectal cancer cell line (modal chromosome number of 45) that is whole-chromosomally stable and expresses wildtype P53. We previously characterized the precise MT dynamics and chromosome instability features of this cell line that is commonly used as a reference model system to study MT function ^22–24^. Results from the NCI-60 panel suggest that ST-401 and NOC inhibited HCT116 cell growth with comparable TGI values of 107 and 141 nM, respectively, as assessed by measuring total protein (**Figure 1b**). However, further analysis of this concentration-dependent response indicated that ST-401 might be more potent than NOC at killing HCT116 cells when tested at lower nanomolar concentrations (**Figure 1e**). Thus, we selected HCT116 cells to study and compare the concentration-dependent antitumor activities of ST-401 and NOC.

### NOC profoundly disrupts the mitotic spindle and mitosis, whereas ST-401 predominantly leaves mitotic MT polymer intact, and this enables mitotic exit

It is well-known that NOC profoundly disrupts the mitotic spindle and arrests cells in mitosis prior to anaphase^25^. To determine the effects of ST-401 on the mitotic spindle and cell cycle arrest in more mechanistic details, we evaluated the appearance of tubulin polymer in fixed HCT-116 cells treated with ST-401 as compared to NOC, and we scored the timing of cell division and cell death of treated cells by live microscopic analysis. As expected, a 4h treatment with NOC (50-200 nM) profoundly disrupted spindle morphology and induced complete MT disassembly at 200 nM (**Figure 2a**). By contrast, mitotic spindles remained provisionally intact during mitosis in HCT116 cells treated with ST-401 (50-200 nM) (**Figure 2a**). Note that ST-401-treated HCT116 cells exhibited both congression defects in prometaphase and lagging chromosomes in anaphase at 50 and 100 nM, as well as multiple spindle poles at 200 nM (**Figure 2a**, arrows). Thus, while HCT116 cells under control conditions exit mitosis in ≈ 40 min, HCT116 cells treated with NOC (100 nM) and ST-401 (100 nM) remained in mitotic arrest for approximately 10-20X longer (**Figure 2b**). **Figure 2b** also shows that ST-401-treated HCT116 cells exhibited congression defects and chromosome missegregation during anaphase that led to the formation of micronuclei (see arrow). Of note, many ST-401-treated cells exited mitosis despite apparent disruption of the mitotic spindle and lagging chromosomes identified during anaphase (**Figures 2c**). Thus, a 24 h treatment with NOC increased the mitotic index starting at 10 nM, whereas treatment with ST-401 induced an increase in mitotic index at 100-300 nM. We also quantified the ability of HCT116 cells to exit mitosis in the presence of ST-401 or NOC (**Figure 2d**). We then determined if the treated HCT116 cells exiting mitosis showed defects during anaphase. Specifically, abnormal anaphase was classified as cells presenting any of these phenotypes: chromosome missegregation, multipolar chromosome segregation, and mitotic exit without cytokinesis. Treatment with ST-401 reduced the number of HCT116 cells undergoing normal mitosis in a concentration dependent manner, with almost no effect at 50 nM and disruption of normal mitosis in 85% of cells at 200 nM (**Figure 2e-f**). By contrast, NOC reduced the number of cells that undergo normal mitosis by 92-99% at all concentrations (**Figure 2e-f**). Accordingly, **Figure 2g** shows that most ST-401-treated cells successfully exited mitosis at all concentrations, and NOC treatment reduced the number of cells that were able to exit mitosis (i.e., they died in mitosis). These data enable two conclusions: 1) between 20-40% of cells treated with NOC at concentrations from 50-200 nM that enter mitosis die during mitosis while the remaining cells are able to exit; 2) A higher percentage of HCT116 cells treated with ST-401 at 50 nM and 100 nM exit mitosis, and a higher percentage of these cells present a normal mitosis compared to NOC. Thus, HCT116 cells treated with ST-401 are more likely to successfully complete mitosis than those treated with NOC.

**Figure 2:**
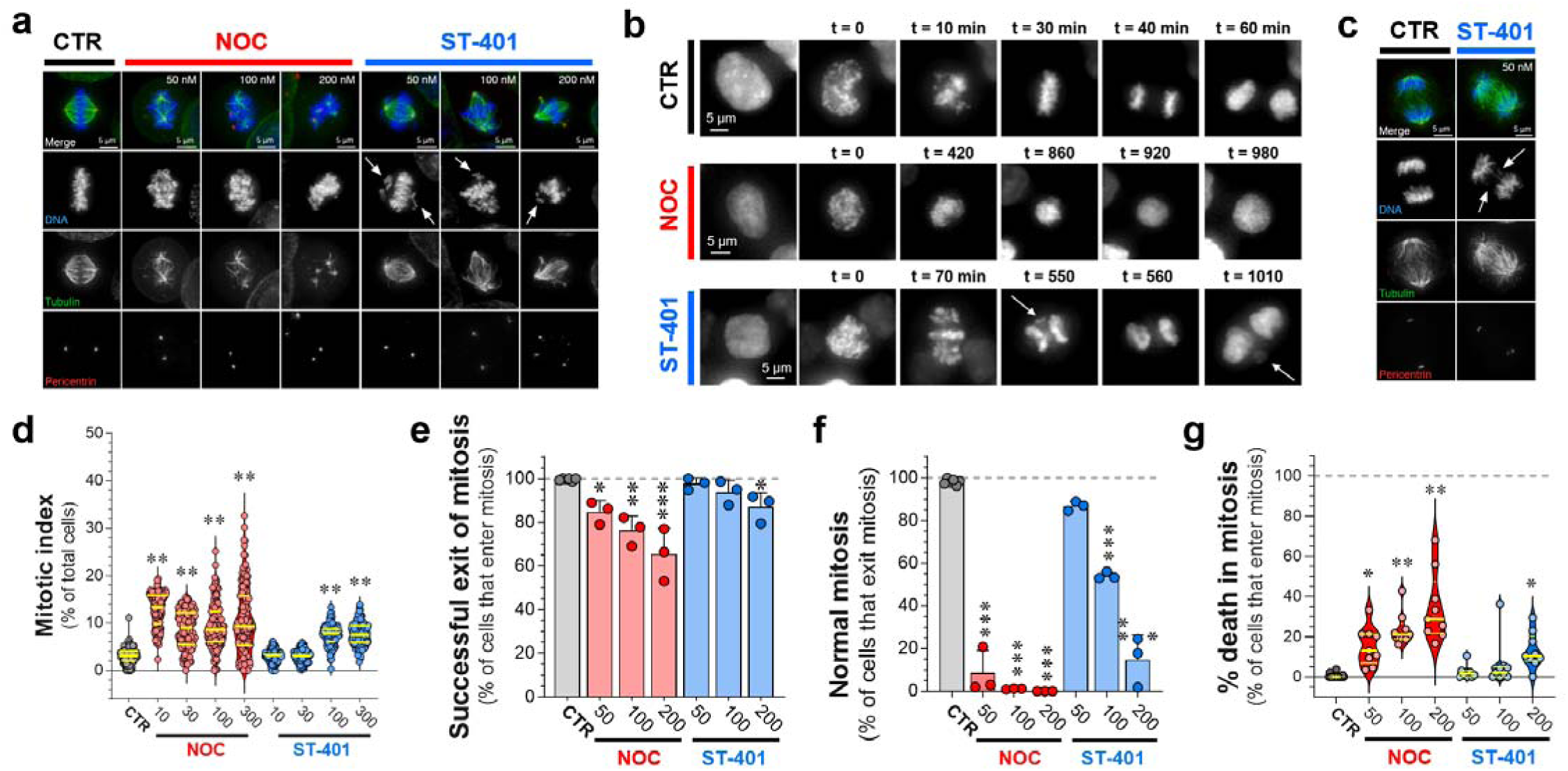
NOC profoundly disrupted the mitotic spindle and mitosis, whereas ST-401 predominantly left mitotic MT polymer intact, which enables mitotic exit. **a)** Immunofluorescence images of HCT116 cells in mitosis after a 4 h treatment with either DMSO, NOC or ST-401. DNA was stained with DAPI (blue), MTs with an anti-tubulin antibody (green) and centrosomes with anti-pericentrin antibody (red). NOC had a more disruptive effect on spindles than ST-401. NOC eliminated a large amount of the MT polymer in the spindles. ST-401 cells showed congression defects (see arrows in DNA channel) and multipolar spindles at 200 nM (pericentrin channel). **b)** HCT116 cells expressing histone H2B fused to mCherry were imaged for 24 h using live fluorescence microscopy in the presence of either vehicle or drugs (see Methods). The nuclear envelope breakdown to anaphase (or DNA decondensation) timing for CTR cells occurred around 40 min, while NOC and ST required several hours to exit mitosis. The NOC-treated HCT116 cell in this example exited mitosis without cytokinesis (just DNA decondensation without cell division, 920 min timepoint). The ST-401-treated HCT116 cells in this example presented congression problems (70 min time point) and lagging chromosomes in anaphase that become micronuclei (arrows in 550 min and 1010 min panels). **c)** Immunofluorescence image of mitotic cells in anaphase after DMSO or ST-401 treatments. The immunofluorescence was performed as in panel a. ST-401 cells presented lagging chromosomes during anaphase (arrows). **d)** Immunofluorescence of cells treated with DMSO, NOC or ST-401 for 24 h were analyzed to calculate the mitotic index (percentage of mitotic cells in a population). DAPI, anti-tubulin antibody and anti-pericentrin antibody were used to detect mitotic structures. We used CellProfiler to identify cells and its machine learning companion CellProfiler Analyst to quantify mitotic cells. NOC yielded a higher mitotic index at all concentrations after 24 h, while ST-401 induced such responses at 100 and 300 nM. For every condition, we recorded 50 random fields with a 10X objective and an 18 mm diameter coverslip. Every spot in the graph corresponds to a field of view, and 3 independent experiments were performed and analyzed. **e**) HCT116 cells tagged with H2B-mCherry were imaged for 24 h in the presence of either DMSO, NOC or ST-401. Most ST-401 treated cells exit mitosis, while a higher percentage of cells in NOC die during mitosis. Data are shown as mean ± s.e.m. of three independent experiments where a total of 2,075 mitotic cells for ST-401 and 3,317 mitotic cells for NOC were analyzed. **f)** Of those cells that exit mitosis and exhibit feature of normal mitosis in the presence of NOC and ST-401 shown in (e), ST-401 treatment of HCT116 cells at increasing concentrations resulted in a higher percentage of cells that exit mitosis without major defects as compared to NOC. **g)** Of those cells that enter mitosis, NOC caused a higher percentage of cell death in mitosis at all concentrations. Statistics (d-g): ***P<0.001, **P<0.01 and *P<0.05 significantly different from corresponding CTR (Two-way ANOVA followed by Dunnett’s). For each, n=3 independent experiments.

### Differential cell cycle responses: NOC treatment resulted in the development of polyploid cells while cells treated with ST-401 avoided this fate

To determine if NOC and ST-401 differentially affected the cell cycle, we analyzed the DNA content of HCT116 cells by flow cytometry using DAPI. As expected, short term treatment (i.e., 4 h) with NOC induced a stronger arrest in G2/M compared to ST-401; for example, NOC (100 nM) increased the number of HCT116 cells in G2/M by 2-fold compared to control, while ST-401 (100 nM) increased the number of HCT116 cells in G2/M by 1.5-fold compared to control (**Figure 3a-b** and Supplementary Figure S2 for quantifications). The differential impact of NOC and ST-401 on the cell cycle was further amplified 24 h after treatment; for example, NOC (100 nM) reduced cells in G1 phase by 15-fold and reduced cells in S phase by 5-fold as compared to CTR; treatment with ST-401 (100 nM) did not affect the number of cells in G1 and S phases compared to CTR (**Figure 3c-f**).

**Figure 3:**
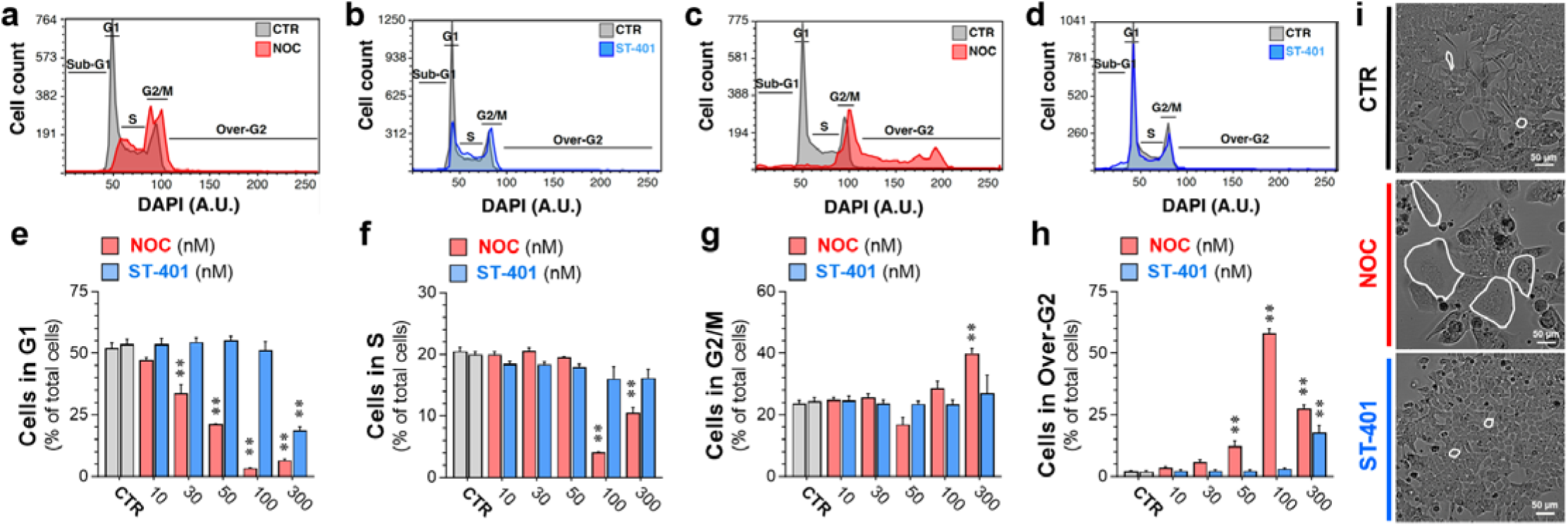
Differential cell cycle responses: NOC treatment resulted in the development of polyploid cells while ST-401 treatment did not. **a-d)** Representative cell cycle plots of HCT116 cells treated with NOC (100 nM) and ST-401 (100 nM) for 4 h (a-b) or for 24 h (c-d). NOC increased the number of cells in G2/M after 4 h and in Over-G2 after 24 h, indicating the formation of polyploid cells, while ST-401-treated HCT116 cells underwent an unperturbed cell cycle (b). **e-g)** Quantification of the number of HCT116 cells in each cell cycle phase when treated with DMSO (CTR) or with increasing concentrations of NOC and ST-401 for 24 h. NOC induced mitotic arrest and increased the number of polyploid cells at concentrations >50 nM, whereas ST-401 induced this response only at 300 nM. Statistics (e-g): Data are expressed as mean ± s.e.m. of n=3 independent experiments. **P<0.01 significantly different from corresponding CTR (Two-way ANOVA followed by Dunnett’s). **i)** Representative bright field images of cells treated with NOC and ST-401 for 5 days. NOC induced the formation of large polyploid cells at lower concentrations.

The increase in the number of cells presenting abnormal anaphases in NOC-treated cells compared to ST-401-treated cells suggest different post-mitotic fates (see Figure 2b). As previously reported, NOC-treated cells that exited mitosis without cytokinesis became polyploid as demonstrated by the increase in the number of cells in the over-G2 phase (e.g., from 2% of total cells in CTR conditions to 58% when treated with NOC (100 nM)) (**Figure 3c&g**) ^26, 27^. In contrast, a 24 h treatment with ST-401 at concentrations up to 100 nM did not generate polyploid cells and at 300 nM only generated 16% polyploid cells (**Figure 3d&g**). Phase contrast microscopy showed that treatment with NOC (100 nM) for 5 days reduced HCT116 cell density and increased the presence of giant cells that resemble PGCC, whereas treatment with ST-401 (100 nM) had no such effect (**Figure 3i**). Accordingly, a 24h treatment with NOC (100 nM) increased cell size as measured by flow cytometry, whereas ST-401 (100 nM) did not (Supplementary Figure S3). Thus, a subset of cells arrested in mitosis in the presence of NOC exited mitosis without completing cell division and persisted as polyploid cells, while ST-401 treatment at the same concentrations avoided this outcome.

### NOC treatment of HCT116 cells induced apoptosis, autophagy, and necrosis, whereas ST-401 treatment avoided these fates and preferentially killed HCT116 cells in interphase

To further investigate the differential cell fate induced by NOC and ST-401, we used live-cell microscopy imaging during a 24 h treatment period to quantify the relative number of cells that died in mitosis versus interphase using the stable cell line HCT116-H2B-mCherry to fluorescently label chromosomes. Specifically, cells dying in mitosis versus interphase were scored based on chromatin condensation and increased DNA fluorescence observed during cell death. Between 2% and 6% of HCT116 cells died under vehicle-treated control conditions (probably due to phototoxicity), with the majority of them dying in interphase and just a few cells dying in mitosis (**Figure 4a**) ^28^. Treatment with NOC at 50-200 nM killed 20-25% of HCT116-H2B-mCherry cells, whereas treatment with ST-401 at the same concentrations killed 8-20% of the cells (**Figure 4a**). Remarkably, ST-401 preferentially killed HCT116 cells in interphase as compared to NOC (**Figure 4a**). For example, 89% of the cells died in interphase with ST-401 at 100 nM compared to 53% of the cells dying in interphase with NOC at 100 nM (**Figure 4a**).

**Figure 4:**
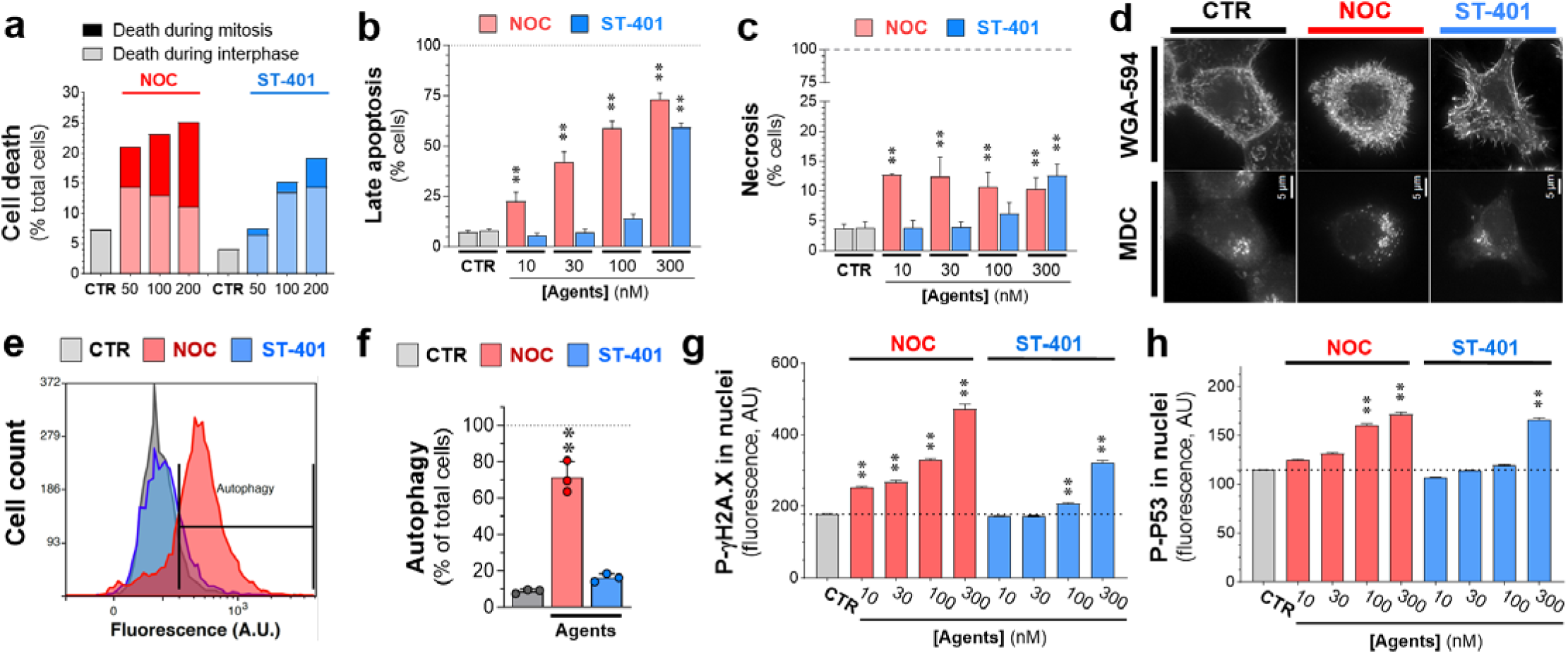
ST-401 kills preferentially in interphase whereas NOC kills preferentially in mitosis and induces apoptosis, autophagy, and necrosis. **a)** Live-cell microscopy imaging of HCT116 cells treated with NOC and ST-401 for 24 h. NOC treatment results in comparable amounts of cell death in mitosis and interphase, while ST-401 treatment results in cells dying more frequently in interphase. Statistics (a). Data are shown as % of total cells from 3 independent experiments (5,500 cells for ST-401 and 7,200 cells for NOC). **b-c)** HCT116 cells treated for 3 days with the MTAs, harvested, and labeled with PI/annexin V to measure apoptosis and necrosis using flow cytometry. NOC dose-dependently and gradually increases apoptosis levels whereas ST-401 induces an all-or-nothing increase in apoptosis at 300 nM. Statistics (b-c): Data are expressed as mean of 3 independent experiments with >10,000 cells. Ordinary one-way ANOVA analysis. **P<0.01 significantly different from corresponding CTR. **d-f)** HCT116 cells stained with monodansylcadaverine (MDC) to detect autophagosomes in HCT116 cells treated with vehicle (CTR), NOC (100 nM) or ST-401 (100 nM) for 24 h. Fluorescence microscopy image of HCT116 cells (d). Flow cytometry histogram of MDC fluorescence (e). Quantification of MDC fluorescence by flow cytometry (f). NOC induces autophagy whereas ST-401 does not. Statistics (e-f): Data are expressed as mean ± s.e.m. of n=3 independent experiments. **P<0.01 significantly different from corresponding CTR (Two-way ANOVA followed by Dunnett’s). **P<0.01 significantly different from corresponding CTR. **g-h)** HCT116 cells were treated vehicle (CTR), NOC (100 nM) or ST-401 (100 nM) for 24 h, immunostained for P-γH2AX and P-P53, imged by fluorescence microscopy and pixel intensity analyzed. NOC increased the levels of both P-H2AX (g) P P53 (h) in the nucleus in a concentration-dependent manner, whereas ST-401 only induced such response at higher concentrations. Statistics (g-h): Data are shown as median with 95% CI for 3 independent experiments where 23,000 and 10,000 total cells were measured for ST-401 and NOC, respectively. Ordinary one-way ANOVA analysis. **P<0.01 significantly different from corresponding CTR.

Based on this finding, we determined whether ST-401 triggered other cell death pathways frequently induced by NOC, such as apoptosis and necrosis. We also checked for the accumulation of autophagosomes that could lead to death by autophagy. To measure apoptosis and necrosis, we treated HCT116 cells with NOC and ST-401 for 1-, 3- and 5-days, and measured annexin-V (**AV**) and propidium iodide (**PI**) staining by flow cytometry. Treatment with NOC and ST-401 for 1-day increased the number of cells in early and late apoptosis. For example, 8% of CTR-treated cells were in late apoptosis and 17% and 13% of cells were in late apoptosis when treated with NOC (100 nM) and ST-401 (100 nM), respectively (Supplementary Figure S4a-b). Neither drug induced necrosis in a 24 h treatment period (Supplementary Figure S4a-b). Substantial differences between ST-401 and NOC emerged after 3 days treatments. NOC induced pronounced late apoptosis (e.g., reaching 59% of cells when treated with NOC at 100 nM) while having little effect on early apoptosis (e.g., reaching only 7.3% of cells when treated with NOC 100 nM) (**Figure 4b** and Supplementary Figure 4Sa-b). By contrast, ST-401 treatment induced late apoptosis only at 300 nM (i.e., 59% of total ST-401-treated cells) (**Figure 4b**). NOC triggered necrosis after 72 h, as shown by an increase of HCT116 cells that stained for PI and not for AV, (e.g., from 4% in CTR-treated cells to 13% in NOC (10 nM)-treated cells) (**Figure 4c**). In contrast, 72 h treatment with ST-401 only triggered necrosis in 13% of HCT116 cells at 300 nM (**Figure 4c**). Thus, while ST-401 is more potent than NOC when measuring TGI inhibition in the NCI-60 cancer cell line panel, it is less potent than NOC at inducing apoptosis and necrosis, suggest an alternative cell death mechanism.

We then surveyed accumulation of autophagosomes labeled by the fluorescent biomarker monodansylcadaverine (**MDC**) ^29^. Autophagy is a survival mechanism that enables cells to degrade damaged cellular components that are no longer needed and can potentially harm cells. However, cells will die when the accumulation of autophagosomes is too high ^30^. Treating HCT116 cells for 24 h with NOC (100 nM) led to accumulation of autophagosomes in 62% of the cells as detected by immunofluorescence (IF) microscopy and quantified by flow cytometry (**Figures 4d-f**). ST-401 (100 nM), however, did not trigger a significant response (**Figures 4d-f**). Prolonged mitotic arrest leads to DNA damage, which triggers the intrinsic apoptosis pathway through TP53. In fact, defects in chromosome segregation and DNA damage during anaphase trigger the phosphorylation and accumulation of phosphorylated P53 (**P-P53**) in nuclei in the subsequent interphase ^31^. **Figures 4g-h** show that NOC increased in a concentration-dependent manner both P-γH2A.X, an index of DNA damage, and P-TP53 levels in nuclei as measured by IF analysis after a 24 h treatment; whereas only the highest level of ST-401 (300 nM) triggers such a response. Together, these results agree with the previous studies showing that NOC (100 nM) kills cancer cells by disrupting mitosis and inducing apoptosis, necrosis, and autophagy, as well as increasing nuclear DNA-damage and P-P53 accumulation. In contrast, lower concentrations of ST-401 do not cause this fate.

### scRNAseq analysis reveals distinct mRNA signatures associated with NOC and ST-401 treatments, including apoptosis, hyperploidy, RNA/protein synthesis and mitochondrial function

The differential antitumor activities triggered by NOC and ST-401 at 100 nM provided an opportunity to distinguish the mRNA signatures associated with death in mitosis versus interphase, as well as the mRNA signatures associated with development of PGCCs, respectively. We performed scRNAseq analysis by single-cell combinatorial indexing RNA-seq (sci-RNA-seq3) of biological replicates to determine the mRNA signatures and cell state of HCT116 cells treated with vehicle (CTR), NOC (100 nM) or ST-401 (100 nM) for either 8 or 24 h (**Figure 5a**) ^32^. In total, we analyzed the single-cell gene expression profiles of 214,165 cells after filtering cells based on both the total number of transcripts and the fraction of mitochondrial reads (Supplementary Figures S5a-d). Quasi-Poisson regression analysis identified 2635 and 3280 genes that were differentially expressed (**DEG**) as a function of NOC or ST-401 exposure at 8 and 24 h, respectively (FDR < 0.05, |normalized β coefficient| > 0.2) (Supplementary Figures S5e-f).

**Figure 5:**
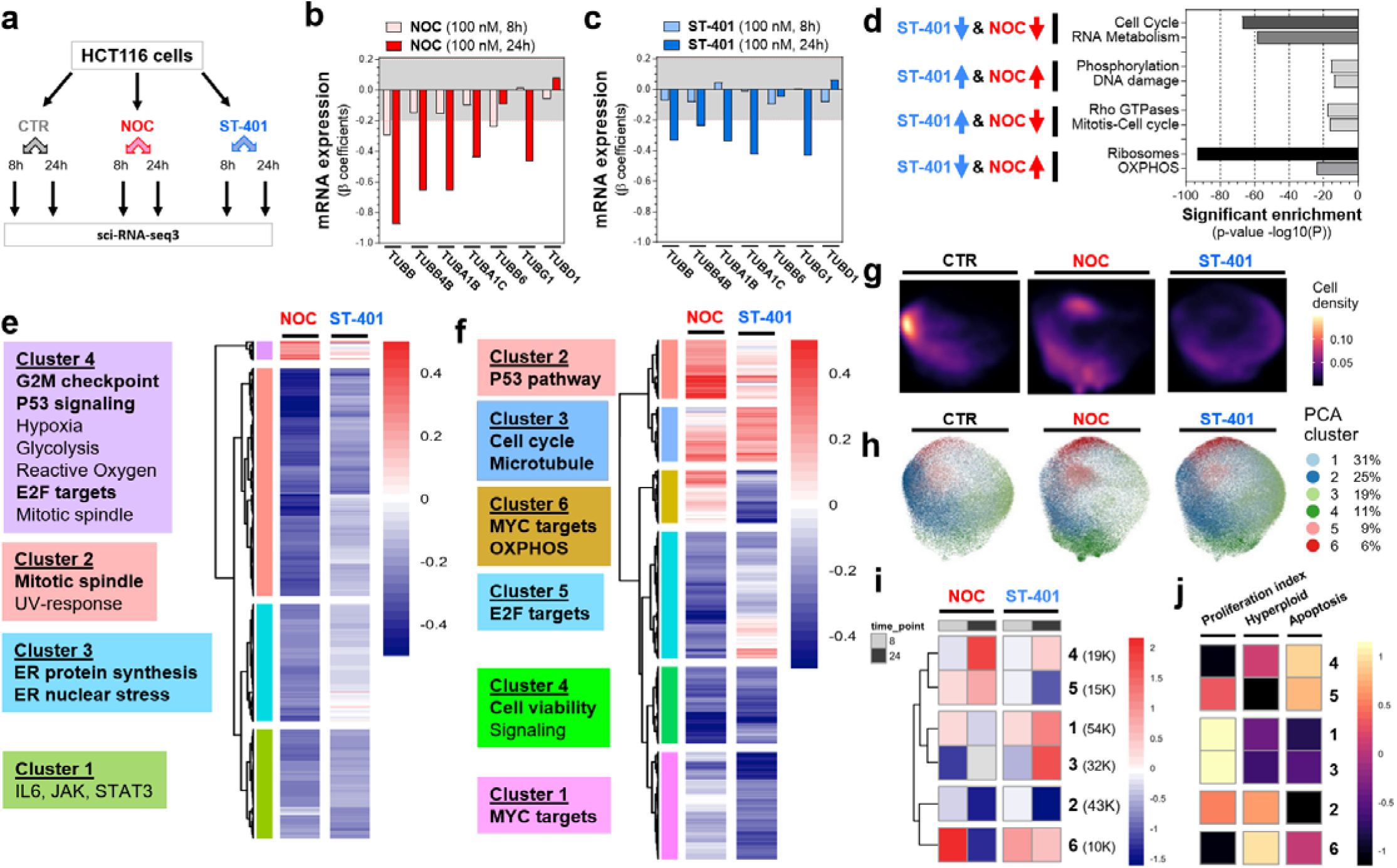
ST-401 and NOC treatment induce differential gene expression and cell states. **a)** Chart depicting the sample structure of our single-cell RNA-seq analysis of NOC and ST-401 exposed HCT116 cells. **b-c)** Change in mRNAs that encode for tubulin subunits following 8 and 24 h treatments with NOC (100 nM) (b) and ST-401 (100 nM) (c). Both X and Y axis represent the normalized effect size (normalized β coefficient) relative to vehicle (CTR) in response to treatment. Statistics (b-c): Gray zone shows filtered for significance (from -0.2 to 0.2). **d)** Bioinformatic analysis using the platform Metascape reveals change in mRNA related to specific cell functions after 24 h treatments with NOC (100 nM) and ST-401 (100 nM) that follow the same and opposite directions. **e-f)** Heatmaps of the union of differentially expressed genes (FDR < 0.05 & abs(β_coefficient_) > 0.2) as a function of NOC or ST-401 treatment for 8 (e) or 24 h (f). Boxes to the right of each heatmap denote significant enriched gene sets for the highlighted gene clusters (FDR < 0.1, hypergeometric test). g. UMAP embedding of single-cell transcriptomes of cells exposed to NOC, ST-401 or vehicle (CTR) for 8 or 24 h faceted by treatment. Cells are colored by cluster and their relative abundance in % of total cells. h. Cell density by condition across the UMAP embedding from g. **g**) Heatmap of the log2 of the odds ratio for the enrichment or depletion of cells across the cell state clusters from CTR treatment. Only statistically significant enrichments are shown (FDR < 0.05, Fisher’s exact test). **h)** Cell density analysis of transcriptomes of cells exposed to NOC, ST-401 or CTR for 24 h. UMAP embedding of single-cell transcriptomes of cells exposed to NOC, ST-401 or vehicle (CTR) for 8 or 24 h. Colors denote Leiden-based community detection cluster assignments in PCA space as calculated using a mutual nearest neighbor approach clustered cells in PCA space with Leiden-based community detection. **i-j)**. Heatmap of the scaled aggregate expression of genes associated with proliferation, hyperploidy and apoptosis for the cell state clusters from (h). Rows (clusters) are ordered as in i. Hyperploid and Apoptosis scores were derived from the MSigDB Chemical and perturbation and the Hallmark gene set collections, respectively.

As expected, α/β-tubulin genes exhibited tubulin autoregulation when treated with MTAs, as shown by both NOC and ST-401 similarly affecting the expression of 6 TUB mRNAs (**Figures 5b-c**) ^33, 34^. Considering that the sensitivity to the ratio of MT polymer to tubulin dimers in the cell controls tubulin mRNA stability, these results suggest that tubulin mRNA stability is similarly affected by NOC and ST-401; and that tubulin autoregulation is unlikely to mediate the differential antitumor activity of NOC and ST-401. First, we used the bioinformatic platform Metascape to identify gene sets associated with biological pathways that are enriched within different modules (i.e., clusters) of differentially expressed genes following 24 h treatments with NOC and ST-401 (Supplementary Figure S5g-h). Thus, **Figure 5d** shows that both NOC and ST-401 induced a robust down-regulation of mRNAs related to the cell cycle and to RNA metabolism, as well as both induced a mild up-regulation in genes related to DNA damage and protein phosphorylation. Notably, ST-401 induced a pronounced down-regulation of mRNAs related to ribosomal assembly and function (ribosomal proteins) and protein translation, as well as those related to mitochondrial OXPHOS and function. In contrast NOC induced an opposite response (**Figure 5d**).

We then used hierarchical clustering to group genes with similar dynamics across conditions. Our analysis showed that 8 h treatments with NOC and ST-401 led largely to the downregulation of gene expression, whereas 24 h treatments resulted in more complex changes (**Figure 5e-f**). We confirmed that treatment of HCT116 cells for 8 h with NOC led to the upregulation of gene clusters related to G2/M check point regulation, TP53 signaling and E2F targets, whereas ST-401 only mildly changed the expression of these gene clusters (**Figure 5e**). The differential impact of NOC and ST-401 on gene clusters was further amplified 24 h after treatment. Specifically, **Figure 5f** shows that treatment of HCT116 for 24 h with NOC lead to the upregulation of gene clusters related to TP53 signaling (Cluster 2) and revealed a downregulation of EF2 targets (Cluster 5). Furthermore, **Figure 5f** confirmed that treatment of HCT116 cells for 24 h led to the downregulation of the gene cluster related to OXPHOS and revealed a downregulation of MYC targets, many of which are associated with ribosomes and protein synthesis machineries (Clusters 6 and 1) ^35^. Thus, NOC preferentially upregulated genes involved in TP53 signaling, ST-401 preferentially downregulates OXPHOS genes, and NOC and ST-401 differentially regulate genes involved in protein translation.

To determine whether NOC and ST-401 induce distinct dynamic changes across cellular states, we performed dimensionality reduction using Principal Component Analysis (**PCA**), we accounted for batch effects between replicates and clustered cells using Leiden-based community detection, and further reduced dimensions using Uniform Manifold Approximation and Projection (**UMAP**) for visualization ^36^. Analysis of cluster distribution for CTR, NOC and ST-401 clusters confirmed the increased differential distributions between CTR, NOC and ST-401 following 24 h treatments compared to 8h treatments (Supplementary Figure S5i). Thus, **Figure 5g** highlights robust differences in the relative accumulation across cellular states following a 24 h treatment. Based on this premise, we investigated the relative accumulation of cells from our condition across 6 cellular states identified by Leiden community detection as a function of treatment time ^37^. **Figure 5h-j** show the phenotypes induced by NOC and ST-401 that are correlated with cellular states as calculated by aggregate gene expression scores based on genes associated with proliferation, hyperploidy and apoptosis. After 8 h, NOC greatly increased scores for cluster 6 that is characterized by a reduced proliferation index, increased apoptosis and increased hyperploidy scores, a score greatly decreased after 24 h (**Figures 5i-j**). By contrast, ST-401 induced a much milder response (**Figures 5i-j**). Furthermore, a 24 h treatment with NOC increased scores for cluster 4, which is also characterized by a reduced proliferation index, increased apoptosis and increased hyperploidy scores, whereas such response was much milder in ST-401-treated cells (**Figures 5i-j**). Treatment for 24 h with ST-401 decreased scores for clusters 2 and 5 and encompassed genes related to a reduced proliferation index (**Figures 5i-j**). Finally, treatment with NOC and ST-401 changed in opposite directions the scores for clusters 3, which includes genes related to the proliferation index (**Figures 5i-j**). Together, our single-cell transcriptomic analysis reinforces our findings that NOC and ST-401 lead to changes in gene expression related to proliferation, hyperploidy, and apoptosis and suggest differential effects on protein synthesis and mitochondrial function.

### Differential integrated stress responses (ISRs) induced by NOC and ST-401

Our scRNAseq analysis indicated that NOC and ST-401 differentially change the expression of genes related to ribosomal proteins and protein translation. We singled out mRNA encoding for ribosomal proteins and plotted the β-coefficients for the agent terms from the DEG tests distributed in quadrants to assess similar and opposite responses. **Figure 6a-b** confirm that mRNAs encoding for cytoplasmic ribosomal proteins are differentially regulated by NOC and ST-401, emphasizing that ST-401 treatment predominantly downregulated these mRNA after both 8 and 24 h treatments as opposed to NOC, which predominantly upregulated these mRNAs. These opposite responses on the expression of cytoplasmic ribosome mRNAs suggest differential effects on protein synthesis and ISR ^38^. Accordingly, we singled out select mRNA encoding for regulators of protein synthesis and ISR, plotted their β-coefficients from the DEG tests. **Figure 6c-d** show that the mRNA coding for ATF4 (the transcription factor regulating ISR output) was downregulated to a greater extent by ST-401 than NOC, and that mRNAs coding for core molecular components of the ISR (eIF2a, EIF2AK2 = PKR, EIF2AK1 = HRI, EIF2AK3 = PERK and EIF2AK4 = GCN2) were differentially regulated.

**Figure 6:**
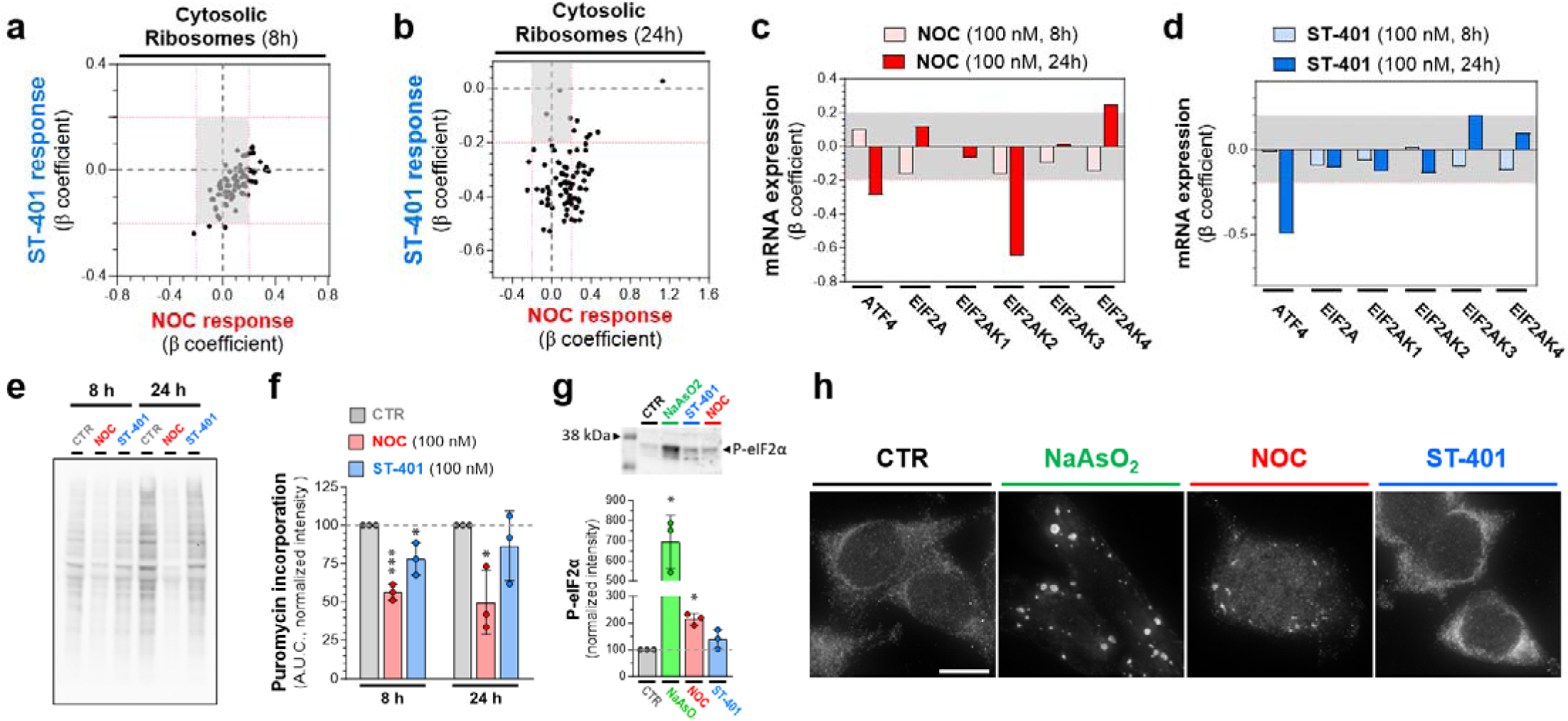
NOC reduced protein synthesis and increased P-eIF2α levels, whereas ST-401 only induced such responses transiently. **a-b)** Changes in the expression of mRNA coding for the cytosolic ribosomal machinery after 8 (a) and 24 h (b) treatment with NOC (100 nM) and ST-401 (100 nM). The x-axis represents the normalized effect size (normalized β coefficient) relative to vehicle (CTR) in response to treatment. **c-d)** Change in the expression of mRNAs involved in the ISR following 8 and 24 h treatments with NOC (00 nM) (c) or ST-401 (100 nM) (d). The x-axis represents the normalized effect size (β coefficient) relative to vehicle (CTR) in response to treatment. Statistics (a-d): Gray zone shows filtered for significance (from -0.2 to 0.2). **e-f)** Western blot analysis of puromycin staining of protein in CTR, NOC (100 nM) and ST-401 (100 nM) treated cells (e) and its quantification normalized by the total amount of protein using Ponceau staining (Supplementary Figure S5a). Statistics (e-f): Data are expressed as means of 3 independent experiments. Ordinary one-way ANOVA analysis (followed by Tuckey). *P<0.05 and **P<0.01 significantly different from corresponding CTR. **g)** Western blot analysis (top) and quantification (bottom) of P-eIF2α levels normalized by total amount of protein using Ponceau staining (Supplementary Figure S5b). Statistics (g): Data are expressed as means of 3 independent experiments. Ordinary one-way ANOVA analysis (followed by Dunnett’s). *P<0.05 and **P<0.01 significantly different from corresponding CTR. **h)** Stress granules labeled with G3BP1 and visualized by fluorescence microscopy. Representative images of HCT116 cells treated for 24 h with vehicle (DMSO 0.1%, CTR), NOC (100 nM) or ST-401 (100 nM) and stained with G3BP1 to visualize stress granules. Positive control: NaAsO_2_ (500 μM). Scale bar = 10 µM.

We used the puromycin assay to measure protein synthesis levels after treatment with NOC (100 nM) and ST-401 (100 nM) and found that NOC reduced protein synthesis by 56% and 50% after 8 and 24 h, respectively while ST-401 reduced protein synthesis by 20% after 8 h and no longer affected protein synthesis after 24 h (**Figure 6e-f**). In line with this result, the levels of P-eIF2α in HCT116 cells treated for 24 h with NOC (100nM) doubled, while ST-401 (100 nM) did not change the phosphorylation of the key regulator of protein synthesis (**Figure 6g**). Increased phosphorylation of eIF2α and development of stress granules play a key role in controlling protein synthesis ^39^. Treatment for 24 h with NOC (100 nM) led to accumulation of stress granules as analyzed by IF using G3BP1 as a marker (**Figure 6h**). **Figure 6h** also shows that the NOC response was less pronounced than the positive control NaAsO_2_ and that ST-401 had no effect on this response ^40^. These results indicate that NOC induces a significant ISR characterized by pronounced and sustained reduction in protein synthesis and pronounced ISR and stress granule formation, whereas ST-401 only induces a transient ISR.

### NOC and ST-401 differentially affect mitochondrial morphology and oxidative phosphorylation (OXPHOS)

Our scRNAseq analysis showed that NOC (100 nM) and ST-401 (100 nM) differentially regulate genes involved in mitochondria OXPHOS function. We singled out mRNA encoding for mitochondrial ribosomal proteins and plotted the β-coefficients for the drug term from the DEG tests distributed in quadrants to assess similar and opposite responses. **Figure 7a-b** show that mRNAs encoding for mitochondrial ribosomal proteins are also differentially regulated by NOC and ST-401: ST-401 treatment predominantly downregulated these mRNA after both 8 and 24 h treatments as opposed to NOC that induced both up and down-regulation of these mRNAs. These differential responses on mitochondrial ribosomal protein mRNAs suggest differential effects on mitochondrial function ^41^.

**Figure 7:**
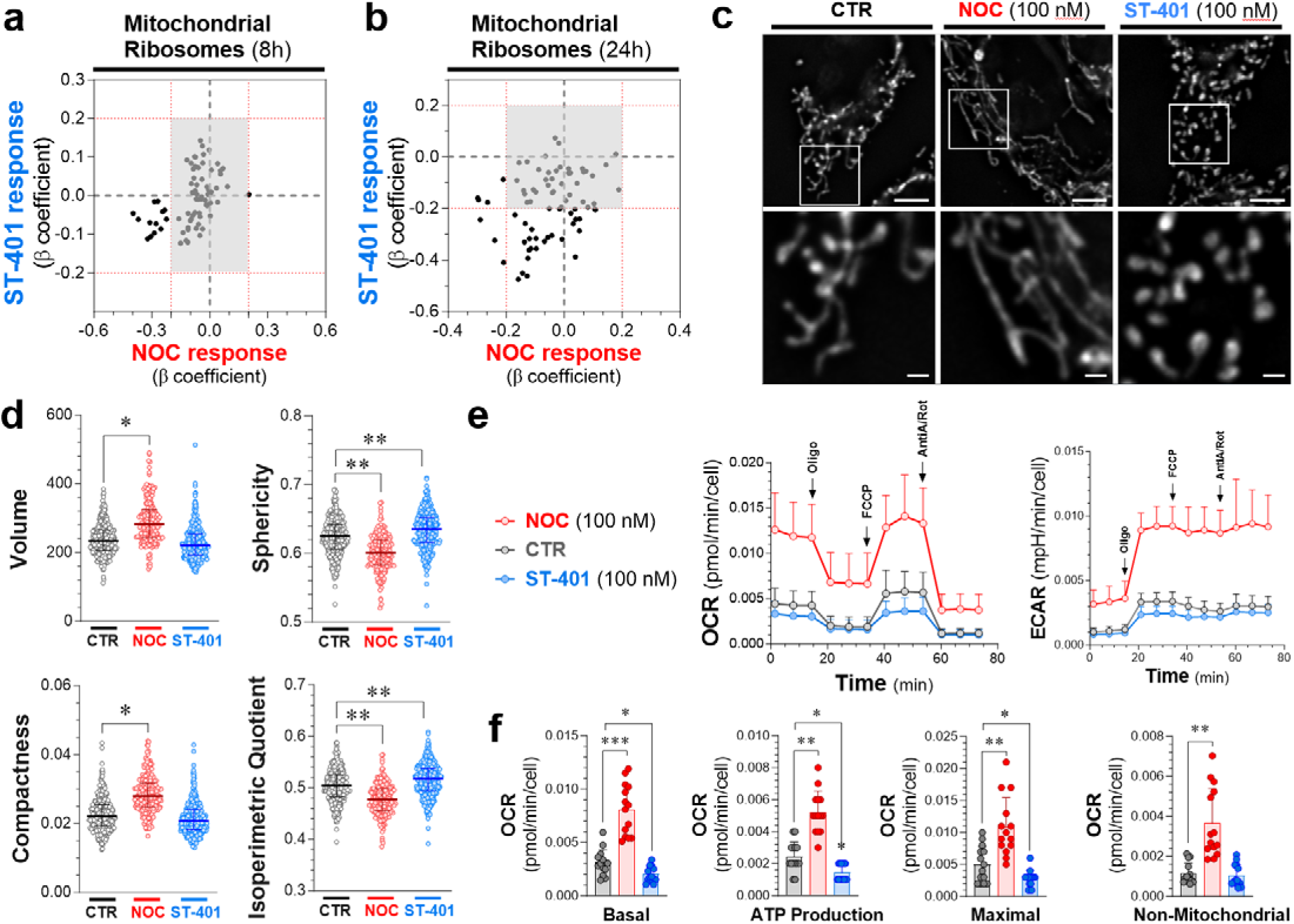
Opposite changes in mitochondrial morphology and OXPHOS induced by NOC and ST-401. **a-b)** Changes in the expression of mRNA coding for mitochondrial ribosomes after 8 (a) and 24 h (b) treatment with NOC (100 nM) or ST-401 (100 nM). The x-axis represents the normalized effect size (normalized β coefficient) relative to vehicle (CTR) in response to treatment. **c-d)** Representative images of HCT116 cells treated for 24 h with vehicle (CTR), NOC (100 nM) or ST-401 (100 nM) and stained with mitoTr cker to visualize mitochondria (c). Unbiased quantification of their volume, sphericity compactness and isoperimetric quotient (d). Statistics (d): Data are shown as mean ± s.e.m. of 3 independent experiments. Ordinary one-way ANOVA analysis resulted in *=p<0.05 and **=p<0.01 compared to CTR. **e-f)** Measure of the oxygen consumption rate (OCR) and of extracellular acidity response (ECAR) of HCT116 treated for 24 h with CTR, NOC (100 nM) or ST-401 (100 nM). Treatment with oligomycin (Olig, 1 μM), FCCP (1 μM), antimycinA/rotenone (1 μM) allowed to resolve the mitochondrial health based on the basal OCR, ATP production, maximal OCR response and non-mitochondrial respiration (f). Statistics (f): Data are shown as mean ± s.e.m. of 5 independent experiments. Ordinary one-way ANOVA analysis resulted in *=p<0.05 and **=p<0.01 compared to CTR.

We analyzed mitochondrial morphology after a 24 h treatment of HCT116 cells with NOC (100 nM) and ST-401 (100 nM) stained with MitoTracker and measuring select parameters of their morphology using MitoMap ^42^. **Figure 7c** shows that NOC treatment induced a pronounced fusion of mitochondria whereas ST-401 induced a pronounced fission. Accordingly, NOC increased the volume and compactness of mitochondria, while reducing their sphericity and isoperimetry quotient (**Figure 7d**). By contrast, ST-401 increased the sphericity and isoperimetric quotient of mitochondria without affecting their volume and compactness (**Figure 7d**). To determine the impact of NOC and ST-401 on OXPHOS, we measured the oxygen consumption rate (**OCR**) and the extracellular acidification rate (**ECAR**) of HCT116 cells treated for 24 h. **Figures 7e-f** show that NOC treatment greatly increased overall OCR, including basal OCR by 2.7-fold and ATP-production by 2.2-fold, as well as several additional OCR parameters (Supplementary Figures S7). By contrast, ST-401 treatment greatly decreased basal OCR by 35% and ATP-production by 41%, as well as several additional OCR parameters (Supplementary Figures S7). NOC induced a comparable increase in ECAR as in its effect on OCR, and ST-401 induced a comparable decrease in ECAR as in its effect on OCR (Supplementary Figures S6). These results show that NOC and ST-401 have opposite effects on the mitochondrial OXPHOS and ECAR response.

## Discussion

The pronounced disruption of MT dynamics during mitosis induced by potent MTAs, such as NOC, delays the correct formation of MT-kinetochore attachments, which elicits the spindle assembly checkpoint and eventually induces cell death following prolonged mitotic arrest. This molecular mechanism, together with spindle defects that remain following MTA treatment, led the field to propose that MTAs primarily kill cancer cell in mitosis ^43^. However, recent studies using drugs that target specific proteins involved in the mitotic machinery (e.g., selective inhibitors of kinases involved in mitotic spindle function) showed limited antitumor activities that never reached the significant antitumor activity achieved by current MTAs in the clinic ^44^. We discovered that ST-401, a milder inhibitor of MT assembly than NOC, induces cell death in interphase at low nanomolar concentrations and cell death in mitosis at higher concentration. This antitumor activity is not mimicked by NOC, suggesting that milder inhibition of MT dynamics may hold broader antitumor activity, killing cancer cells at low concentrations in interphase and at high concentrations also in mitosis.

### How do MTAs induce death in mitosis versus exiting mitosis?

Studies that used live microscopy imaging of cancer cell lines showed that cells within any given cell line exhibit multiple fates following prolonged arrest. Specifically, while some cells die in mitosis, others exit mitosis without dividing and return to interphase, and once back in interphase, some cells undergo cell-cycle arrest, others die, and others re-replicate their genomes ^45^. Importantly, these studies showed that cell fate is not necessarily genetically predetermined and that duration of mitotic arrest does not dictate fate and revealed that delaying mitotic exit shifts the fate profile toward death in mitosis ^45^. Thus, the duration of mitotic arrest does not dictate cellular fate. One possibility is that DNA damage slowly accumulates during a protracted mitosis that, in the absence of efficient repair, eventually activates cell death pathways ^45^. Despite this fundamental evidence, it remains unclear whether death in interphase represents a delayed response to damage incurred during mitosis or whether returning to interphase without successfully dividing creates a de novo damage signal. We show that HCT116 cells treated with low nanomolar concentrations of ST-401 were able to exit mitosis. By contrast, this did not occur in HCT116 cells treated with low nanomolar concentrations of NOC, which increased the mitotic index and triggered cell death via apoptosis, necrosis, and autophagy at low nanomolar concentrations. Accordingly, NOC was more potent at increasing P-P53 and P-γH2AX levels and inducing apoptosis than ST-401, and ST-401 only induced such response at higher concentrations. Of those cells that exit mitosis and escape cell death, near to 100% of the NOC-treated cells exhibited chromosomal defects as compared to ST-401. In fact, some cells were able to exit mitosis without cytokinesis. This was confirmed by flow cytometry as a 24 h treatment with NOC (100 nM) resulted in ≈60% of cells developing polyploidy. Thus, our study suggests that potent inhibitors of MT assembly induce death in mitosis, whereas milder inhibitors of MT assembly allow cells to exit mitosis and then kill cancer cells in interphase at low concentrations and kill cancer cells in mitosis at high concentrations.

The low doubling times and mitotic index of most tumors (between 1 and 3%) indicates that therapeutic approaches that also kill cancer cells in interphase likely hold increased efficacy ^2^. The interphase MT network presents a large surface area for protein-protein interactions, and, combined with the innate polarity of MTs, allows the trafficking of proteins and organelles through the cell via motor proteins, such as kinesin and dynein ^46^. In addition to spatially organizing the cellular contents, this machinery also controls cellular function via several signaling pathways ^8^. We show that ST-401 induces cell death in interphase at low nanomolar concentrations, suggesting that HCT-116 cells in G1 phase are susceptible to cell death. Of note, the mechanism described here appears to differ from the mechanism shown when cells that have passed a putative "microtubule sensitivity checkpoint" mechanism in late G1/early S-phase continue to cycle and die following mitotic arrest ^47^. Thus, the fate profiles of cancer cells can change dramatically at different drug concentrations. A striking difference exists in death response triggered by increasing concentrations of ST-401, vincristine and eribulin. The latter two MTAs bind to adjacent binding sites at the interface of α/β tubulin heterodimers (the *vinca* and *eribulin* sites), and inhibit MT assembly, disrupt mitotic spindles and kill cells in G2/M at lower (< 30 nM) and higher concentrations (> 100 nM) and induce death in G1 phase through a P-P53-dependent mechanism ^8, 47, 48^. Thus, while ST-401 kills cancer cells in interphase at lower concentrations and in mitosis at higher concentrations, vincristine does the opposite, killing cancers cells in mitosis at low concentration and in interphase at high concentration ^48^. These results agree with our NCI-60 Compare analysis showing poor correlations between ST-401 and vincristine’s antitumor activities.

Earlier studies did not determine whether the postmitotic response is a delayed reaction to damage incurred during mitosis or whether returning to interphase without successfully dividing creates a de novo damage signal. In fact, it seems unlikely that such brief delay is sufficient for damage signals to accumulate to a level high enough to trigger a response, strongly suggesting that there is indeed a separate postmitotic response associated with death in interphase that kill PPGCs. Whether other MTAs, such as vincristine and eribulin, induce similar cellular responses involving ISR and disruption of mitochondria in interphase as ST-401, and whether such response varies depending on the type of cancer cells, remains to be determined.

Chromosome segregation during mitosis is a highly regulated process carried out by the mitotic spindle, a multiproteic machinery combined with dynamic MTs ^49^. Thus, proper tuning of MT dynamics within the spindle is essential to avoid the mis-segregation of chromosomes during anaphase and polyploidy. Several outcomes have been described that follow prolonged arrest. While some cells appear to die in mitosis, other cells exit mitosis without dividing and return to interphase. Once back in interphase, some lines undergo cell-cycle arrest, others die, and others re-replicate their genomes, i.e., endocycle, and continue to proliferate ^50^. Cells that exit mitosis with low chromosomal instability (**CIN**) continue through the cell cycle without forming 4n cells and PGCC. However, as these cells undergo more cell cycles, they fail to split chromosomes between daughter cells, accumulate CIN over several cell cycles and eventually die in mitosis or generate PGCC, which have been implicated in the development of cancer and drug resistance ^51, 52^. Accordingly, live tracking of cell fate experiments revealed a remarkable detrimental side-effect of escaping mitotic arrest. Specifically, some mitotically arrested cells that fail cytokinesis can still exit mitosis through mitotic slippage and become PGCCs or cancer-stem-cells implicated in malignancy development, acquisition of metastatic characteristics and drug resistance ^11, 13, 14^. Possibly, the strong disruption of mitosis triggered by NOC that eliminates most MT polymer might favor the selection of cells that carry the ability to replicate without a spindle, a hallmark of PGCCs ^50^. By contrast, ST-401 triggers low levels of CIN and thus does not kill cells in mitosis. Together, these studies raised the possibility that killing cells by mitotic arrest is inefficient and killing cancer cells in interphase might be more efficient ^2, 9, 44^. In fact, such an approach would be quite valuable as PGCCs can persist and even continue to grow by amitotic budding, facilitating their invasive potential, and thus become cancer stem cells that initiate dedifferentiation in response to therapy-induced stress ^12, 15^.

Our scRNAseq analysis unraveled potential mechanisms associated with cell death in interphase that involve the ISR and impaired mitochondrial function. Cells respond to different types of stress by inhibition of protein synthesis and subsequent assembly of stress granules, which are cytoplasmic aggregates that contain stalled translation preinitiation complexes ^53^. Specifically, the rate of protein synthesis in cells is tightly regulated, including through eIF2α. For example, in response to various forms of stress, cells reduce global translation, by which they prevent further protein damage, reallocate their resources to repair processes, and ensure cellular survival ^54^. Thus, the packaging of cytoplasmic mRNA into discrete RNA granules regulates gene expression by delaying the translation of specific transcripts. Following translation, however, newly inactivated mRNAs released from polysomes can also be packaged into dynamic, transient structures known as stress granules. Significantly, protein synthesis is essential for the formation and growth of several types of cancer and often dispensable for non-malignant tissue homeostasis. Thus, such tumors are often therapeutically vulnerable to protein synthesis inhibitors ^55^. Autophagy is a regulated process for the degradation of damaged organelles or proteins activated by different types of stresses ^56^. The damaged organelles or proteins are stored in autophagosomes that fuse to lysosomes for the elimination of those harmful components. Thus, autophagy allows cells to survive mild stressful conditions. We found that NOC induces prolonged ISR and autophagy, whereas ST-401 induces a transient ISR that does not result in the accumulation of stress granules and induction of autophagy after a 24 h treatment. Thus, the cell death mechanism induced by ST-401 in interphase does not involve autophagy despite a transient ISR.

### Opposite changes in mitochondria morphology and OXPHOS

Converging lines of evidence indicate that mitochondria play a key role in the biological embedding of adversity, including linking the consequence of stress exposure to changes in mitochondrial structure and function ^53^. This molecular link and its reprogramming are now recognized hallmarks of cancer as recent studies have uncovered remarkable flexibility in the specific pathways activated by tumor cells to support these key functions ^57^. Thus, tumors reprogram pathways of nutrient acquisition and metabolism to meet the bioenergetic, biosynthetic, and redox demands of malignant cells ^58^. Impairment of mitochondrial function by the ISR can lead to several problems for cancer cells, including reduced cell growth and increased cell death linked to reduced ATP levels. We found that NOC and ST-401 induce opposite responses on mitochondria morphology and OXPHOS: NOC induced mitochondrial fusion and increased mitochondrial volume and increased OXPHOS, whereas ST-401 induced mitochondrial fission without a changed volume and decreased OXPHOS. The mitochondrial phenotype induced by NOC is likely associated with increase superoxide production and contributes to the killing of HCT-116 cells by apoptosis, necrosis, and autophagy ^59^. The mitochondrial phenotype induced by ST-401 characterized by a reduction in energy metabolism is likely to contribute to the killing of HCT-116 cells in interphase. Importantly, while most mitochondrial proteins are synthesized in the cytoplasm, select mitochondrial proteins, such as mitochondrial ribosomal proteins (**MRPs**), are synthesized in the cytosol and transported to mitochondria ^60^. In fact, MRPs are elevated in select cancers and several approaches have been developed to target MRPs for the treatment of cancer ^60, 61^. Considering that NOC and ST-401 differentially affect MRP expression, these results suggest that NOC and ST-401 treatment might also directly affect mitochondrial function independently of IRS, including by differentially affecting MRP expression directly. Thus, our results suggest that death in interphase induced by mild inhibition of MT assembly in interphase is likely to involve significant reduction in energy supply.

<u>In conclusion</u>, our study unravels a molecular mechanism that mediates the death of cancer cells in interphase and that it involves impaired mitochondrial function characterized by fission and reduction in energy metabolism. Mild inhibition of MT assembly is likely to induce broad antitumor activity, killing cancer cells in interphase at low concentrations and in mitosis at high concentrations as compared to strong inhibition of MT assembly that principally kills cancer cells in mitosis and might lead to the development of PGCC.

### Funding, Disclosure and Acknowledgments

This work was supported by the National Institutes of Health [Grants NS106924 and CA244213 to N.S, GM069429 to J.J.V. and DP2ES032761 to YS). J.J.V. and Y.S. were also supported by the University of Washington Royalty Research Fund. NS is employed by Stella Consulting LLC. The terms of this arrangement have been reviewed and approved by the University of Washington in accordance with its policies governing outside work and financial conflicts of interest in research. We thank Dr. Cailyn Spurrell from the Brotman Baty Institute for Precision Medicine, University of Washington Seattle, for help with processing the scRNAseq samples, Drs. Mark Kunkel and Ernest Hamel from the National Cancer Institute Developmental Therapeutics Program in Bethesda for help with analyzing NCI-60 cancer panel data, and Dr. Linda Wordeman for insightful discussions and constructive edits. We also thank the UW Pathology Flow Cytometry Core Facility and its manager Dr. Aurelio Silvestroni.

## Methods

### Statistical analysis

GraphPad Prism 9: curve fit (three parameters), Multiple comparison (ANOVA followed by DUNETT’s or TUCKEY’s).

### NCI-60 screen and COMPARE

The NCI-60 cell line screen is a service available to the cancer research community. The screening process, data processing, and the COMPARE algorithm are described: https://dtp.cancer.gov/databases_tools/docs/compare/compare_methodology.htm Briefly, cells are established in 96 well plates for 24 h, exposed to test compounds across a range of concentrations for 48 h and viability determined by sulforhodamine B dye exclusion assay. Data sets are created for the concentrations at which compounds caused 50% growth inhibition (GI50), total growth inhibition (TGI) or 50% cell kill (LC50). The compare algorithm works with the sets of data, returning either every possible pairwise correlation within a defined set of data (grid COMPARE) or uses a data set (seed) to determine the highest correlations within a defined set (targets) of data (standard COMPARE).

### Cell Culture

Colon cancer cell line-HCT 116 was purchased from ATCC and cultured in RPMI media with 10% FBS and 1% penicillin/streptomycin. Cells were maintained at 37 °C and 5% CO_2_. When cells were 70-80% confluent they were subcultured using trypsin-EDTA.

### Cell Fixation and Immunofluorescence

HCT116 cells were treated with NOC or ST-401 and were fixed after 4 h using 2% paraformaldehyde in methanol at -20 °C for 10 min. Tubulin and centrosomes were labeled using anti-tubulin monoclonal DM1-alpha (1:500, Sigma) or anti-pericentrin (1:500, Abcam) antibodies applied overnight at 4 °C. Alexa-fluor 488 and 647 secondary antibodies against mouse and rabbit IgG, respectively, were used to detect the anti-tubulin and anti-pericentrin antibodies. Coverslips were mounted using ProLong® Diamond containing DAPI (Molecular Probes, Eugene, OR) and cured for 24 h. Fluorescent images were collected as 0.5-μm Z-stacks on a Deltavision microscope system (Applied Precision/GEHealthcare, Issaquah, WA) using a 60x 1.42 NA lens (Olympus, Tokyo, Japan). Images were deconvolved using SoftWorx 5.0 (Applied Precision/GEHealthcare), and representative images are presented as a flat Z-projection. Figure 4 c,d: HCT116 cells were treated with NOC or ST-401 and were fixed with 4% paraformaldehyde in PBS at 37 °C for 15 min and permeabilized in 0.5% Triton X-100 for 5 min at room temperature. P-p53 and P-γH2AX were labeled using a mouse anti-P-p53 (1:250, Phospho-p53 (Ser15) (16G8), Cell Signaling Technology) or a rabbit anti-P-H2AX (1:250, Phospho-Histone γH2A.X (Ser139) (20E3), Cell Signaling Technology) applied overnight at 4 °C. Alexa-fluor 488 and 568 secondary antibodies against mouse and rabbit IgG, respectively, were used to detect the anti-P-p53 and anti-P-γH2AX antibodies. Coverslips were mounted using ProLong® Diamond containing DAPI (Molecular Probes, Eugene, OR) and cured for 24 h. Fluorescent images of random fields of the coverslips were collected using a GE InCell 2500. Automatic identification of nuclei and fluorescence intensity was performed using the free software CellProfiler.

### Live-cell Imaging

For live-cell imaging, a stable cell line HCT116-mCherry-H2B cells were cultured in glass-bottom 24 multi-well plates from Ibidi. Cells were switched to 37 °C in a CO_2_ independent media (10% FBS) with either DMSO, NOC or ST-401. Live-cell images were acquired using a GE InCell 2500 system with a 20X lens. Time-lapse images were collected as 15 µm Z-stack series, with 3 µm Z-spacing, and with 10 min time intervals for 24 h. Images were deconvolved using the InCell software and processed with Fiji. All quantification was done manually using Fiji.

### Cell Cycle Experiments

HCT116 cells were treated with either DMSO, NOC or ST-401 at different times. After drug treatment, cells were collected using trypsin and centrifuged at 1,000 r.p.m. for 10 min. The cell pellets were resuspended in 400 µL of a 10 µg/mL DAPI, 0.1% NP40 Tris buffer. Cells were processed by flow cytometry, and cell cycle data was analyzed using FlowLogic software.

### Apoptosis/necrosis Experiments

HCT116 cells were treated with either DMSO, NOC or ST-401 at different times. After drug treatment, cells were collected using trypsin and centrifuged at 1,000 r.p.m. for 10 min. Cells were treated with a necrosis/apoptosis kit from ThermoFisher containing PI and Annexin V Alexa Fluor™ 488 according to the manufacturer specifications. In short, the cell pellets were resuspended in 100 µL of annexin binding buffer, followed by treatment with 5 µL Alexa fluor annexin V-488 and 1 µL of 100 µg/mL PI. After 15 min at room temperature, 400 µL of buffer was added to the mix. Cells were processed by flow cytometry, and apoptosis/necrosis data was analyzed using FlowLogic software.

### PI and Annexin V Staining

PI is an impermeable DNA-binding dye that enters cells when the cell membrane is compromised. PI, when excited, emits light at around 620 nm (orange/red) and it is routinely used in flow cytometry as a necrosis marker. Annexin V is a phosphatidylserine-binding protein that is used as an apoptosis marker, and when fused to AlexaFluor 488, emits light at around 525 nm (green).

### Puromycin incorporation

HCT116 cells were treated with either DMSO, NOC or ST-401 at different times. After drug treatment, cells were treated with 1 µM Puromycin (Thermo Scientific Chemicals) at 37 °C for 30 min. After Puromycin treatment cells were collected using trypsin and centrifuged at 1,000 r.p.m. for 10 min. The cell pellets were resuspended in 100 µL of RIPA buffer with protease inhibitors and incubated in a rocking platform shaker at 4 °C for 60 min. After it, the samples were centrifugated at 12,000 r.p.m. at 4 °C for 30 min. The supernatant was used for a western-blot experiment using a mouse-anti-Puromycin antibody (1:5,000, Millipore MABE343). To detect primary Ab, we incubated samples with donkey-anti-mouse Ab fused to HRP (1:5,000 Jackson ImmunoResearch 715-036-151). Pierce ECL Western Blotting Substrate was used for HRP detection. Signals were detected with a Bio-Rad ChemiDoc MP Imaging System.

### P-eIF2a experiments

HCT116 cells were treated with either DMSO, NOC or ST-401 at different times. After drug treatment, one control sample was treated with 500 µM NaAsO_2_ for 1 h at 37 °C. After this treatment cells were collected using trypsin and centrifuged at 1,000 r.p.m. for 10 min. The cell pellets were resuspended in 100 µL of RIPA buffer with protease inhibitors and incubated in a rocking platform shaker at 4 °C for 60 min. After it, the samples were centrifugated at 12,000 r.p.m. at 4 °C for 30 minutes. The supernatant was used for a western-blot experiment using a mouse-anti-Phospho-eIF2α (Ser 51, 1:5,000, Proteintech 68023-1). To detect primary Ab, we used a donkey-anti-mouse Ab fused to HRP (1:5,000 Jackson ImmunoResearch 715-036-151). Pierce ECL Western Blotting Substrate was used for HRP detection. Signals were detected with a Bio-Rad ChemiDoc MP Imaging System.

### Autophagy

Flow cytometry: HCT116 cells were treated with either DMSO, NOC or ST-401 for 24 h. After drug treatment, cells were incubated with 50 µM of MDC (Sigma-Aldrich 30432) for 45 min at 4 °C in the dark. After MDC incubation, cells were washed with PBS 3 times, detached using trypsin and centrifuged at 2,000 r.p.m. for 10 min. Cell pellets were washed another 3 times with PBS and resuspended in 1 mL of PBS. Cells were processed by flow cytometry using a BD LSR II Flow Cytometer, and data was analyzed using FlowLogic software. Microscopy: HCT116 cells were treated with either DMSO, NOC or ST-401 for 24 h. After drug treatment, cells were incubated with 50 µM MDC for 15 min at 37 °C in the dark. Cells were then fixed with 4% PFA in PBS at 37 °C for 15 min and permeabilized in 0.5% Triton X-100 for 5 min at room temperature. After a PBS washes, cells were incubated with WGA-594 (Wheat Germ Agglutinin fused to a fluorophore) at RT for 10 min to label the cell membrane. After PBS wash, coverslips were mounted using ProLong® Diamond containing DAPI (Molecular Probes, Eugene, OR) and cured for 24 h. Fluorescent images were collected as 0.5-μm Z-stacks on a Deltavision microscope system (Applied Precision/GEHealthcare, Issaquah, WA) using a 60x 1.42 NA lens (Olympus, Tokyo, Japan). Images were deconvolved using SoftWorx 5.0 (Applied Precision/GEHealthcare), and representative images are presented as Maximum intensity Z-projection.

### G3BP1

HCT116 cells were treated with either DMSO, NOC or ST-401 for 24 h. After drug treatment, a CTR sample was treated with 500 µM NaAsO_2_ for 1 h at 37 °C. Cells were then fixed with 4% PFA in PBS at 37 °C for 15 min and permeabilized in 0.5% Triton X-100 for 5 min at room temperature. G3BP1 was labeled using a mouse-anti-G3BP1 (1:250, Proteintech, 66486-1) applied overnight at 4 °C. A donkey-anti-mouse Ab Alexa-fluor-488 (1:500) was used to detect the anti-G3BP1 Ab. After PBS wash, coverslips were mounted using ProLong® Diamond containing DAPI (Molecular Probes, Eugene, OR) and cured for 24 h. Fluorescent images were collected as 0.5-μm Z-stacks on a Deltavision microscope system (Applied Precision/GEHealthcare, Issaquah, WA) using a 60x 1.42 NA lens (Olympus, Tokyo, Japan). Images were deconvolved using SoftWorx 5.0 (Applied Precision/GEHealthcare), and representative images are presented as Maximum intensity Z-projection.

### Mitochondria morphology

HCT116 cells were plated in 96-well plates with glass bottom from In Vitro Scientific, USA (IVS-P96-1.5H-N). After treatment for 24 h with DMSO, NOC or ST-401, cells were incubated with 100 nM MitoTracker green FM (Invitrogen M7514) for 45 min at 37 °C. After a media wash, fluorescent images of random fields were collected as 0.3-μm Z-stacks using a GE InCell 2500 with a 60X lens. Images were processed using FiJi and analyzed for mitochondrial morphology using the FiJi plugin MitoLoc (a.k.a.MitoMap).

### Mitochondrial Function

The Cell Mito stress test was employed to assess key metabolic parameters, including Oxygen Consumption Rate (OCR), Extracellular Acidification Rate (ECAR), and ATP production, using the Agilent XFe96 Seahorse analyzer (Agilent Technologies, Wilmington, DE). HCT116 cells (0.75 x 104) were seeded into a 96-well plate, treated with specific compounds, and incubated for 24 h. OCR and ECAR were measured under basal conditions, and, subsequently, 1 µM oligomycin, 1 µM FCCP, and 1 µM antimycin A/rotenone were added to the cells. Relevant parameters, such as basal and maximal respiration, ATP production, proton leak, non-mitochondrial respiration, and spare respiratory capacity, were calculated in accordance ^62^. Data were normalized based on cell numbers determined through microscopy analysis.

## Abbreviations

(**AV**): annexin-V
(**CIN**): chromosomal instability
(**DEG**): differentially expressed
(**ECAR**): extracellular acidification rate
(**ISR**): integrated stress response
(**MTs**): microtubules
(**MTAs**): microtubule targeting agents
(**MRPs**): mitochondrial ribosomal proteins
(**MDC**): monodansylcadaverine
(**NOC**): nocodazole
(**P-P53**): phosphorylated P53
(**OXPHOS**): oxidative phosphorylation
(**OCR**): oxygen consumption rate
(**PGCC**): polyploid giant cancer cells
(**PCA**): Principal Component Analysis
(**PI**): propidium iodide
(**SAC**): spindle assembly checkpoint
(**TGI**): total growth inhibition
(**UMAP**): Uniform Manifold Approximation and Projection

