## Supplemental figures for "Mitosis exit followed by death in interphase prevents the development of polyploid giant cancer cells"

### Supplementary Figures

**Supplementary Figures S1a-b:** $GI_{50}$, TGI and $LC_{50}$ MeanGraph and concentration/response curves for ST-401.

**Supplementary Figure S1c:** NCI60 Mean Graphs – 12 highest correlations between ST-401 and Approved Agents (169 NSC).

**Supplementary Figure S2:** Four h treatment with NOC disrupts the cell cycle of HCT-116: milder response by ST-401.

**Supplementary Figure S3:** Cells exiting mitosis: NOC without cytokinesis and ST-401 with a lagging chromosome.

**Supplementary Figures S4a-c**: Quantification of MDC levels in HCT116 treated cells measured by flow cytometry.

**Supplementary Figure S4d:** NOC trigger apoptosis: milder response by ST-401**.**

**Supplementary Figures S5a-d:** Overview of single-cell RNA-seq sample metrics.

**Supplementary Figures S5e-f:** Summary of differential gene expression analysis of vehicle, NOC and ST-401 exposed cells.

**Supplementary figures S5g-h:** Gene-enrichment analysis at 24h using Metascape.

**Supplementary figure S5i:** Gene-enrichment analysis at 24h using Metascape.

**Supplementary figure S6a-b:** Ponceau Red staining of gels shown in Figure 6.

**Supplementary figure S7:** NOC (100 nM) and ST-401 (100 nM) treatment for 24 h increases and decreases OCR and ECAR, respectively.

**Graphical Abstract:** Diagram depicting ST-401 antitumor activity.

**Supplementary Figures S1a-b:** GI50, TGI and LC50 MeanGraph and concentration/response curves for ST-401: **a)** MeanGraph representation of the concentrations in the NCI60 cell lines for ST-401 to cause GI50 (50% growth inhibition), TGI (total growth inhibition) or LC50 (50% cell kill) in the screen. The midline for each graph is the average concentration for all of the cell lines at that endpoint. Cell lines where are more sensitive than average have bars drawn to the right. **b)** The concentration/response data are also presented. Correlations among the three endpoint patterns were low. The TGI endpoint was used for subsequent work since the TGI endpoint has been found to be a better indicator for mechanisms which target tubulin, and a lower correlation limit of 0.6 was used for significance. (ref: Hamel)

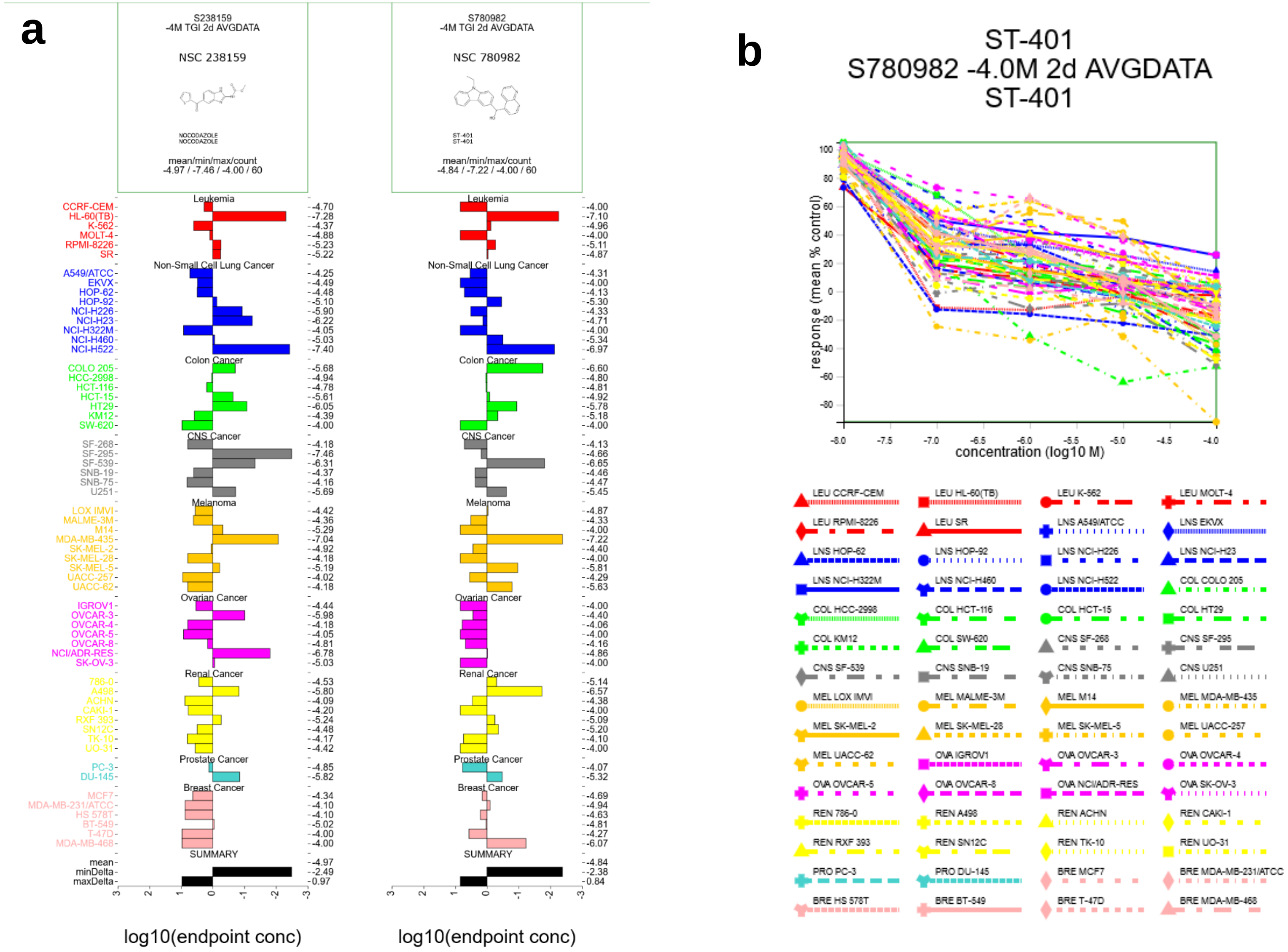

**Supplementary Figure S1c:** NCI60 Mean Graphs – 12 highest correlations between ST-401 and Approved Agents (169 NSC): MeanGraph showing the TGI patterns for ST-401 and the most-highly correlated TGI patterns in the NCI60 set of Approved Agents, ordered by decreasing correlation.

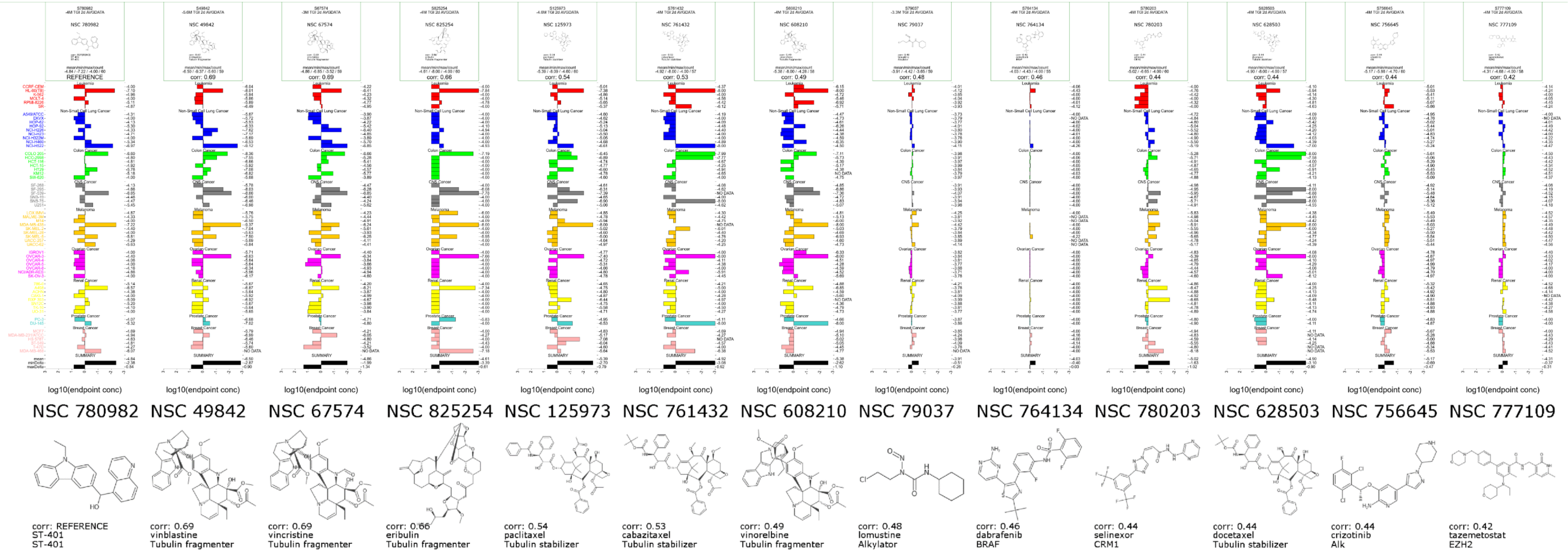

**Supplementary Figure S2:** Four h treatment with NOC disrupts the cell cycle of HCT-116: milder response by ST-401. Data were analyzed by ANOVA multi-comparison test followed by DUNETT's post test. N = 3 independent experiments. * = P < 0.0332, ** = P < 0.021, *** = P < 0.0002 and **** = P < 0.0001 different from Vehicle.

| 4 h | Sub-G1 | | | G1 | | | S | | | G2/M | | | Over-G2 | | |
|---|---|---|---|---|---|---|---|---|---|---|---|---|---|---|---|
| | Mean | SD | p | Mean | SD | p | Mean | SD | p | Mean | SD | p | Mean | SD | p |
| **CTR** | 0.9 | 0.2 | | 51.2 | 3.9 | | 21.1 | 2.1 | | 23.7 | 1.8 | | 3.1 | 1.4 | |
| 10nM ST | 1.3 | 1.0 | ns | 49.2 | 4.9 | ns | 22.2 | 2.6 | ns | 23.2 | 2.7 | ns | 4.0 | 1.4 | ns |
| 30nM ST | 0.9 | 0.2 | ns | 47.5 | 4.6 | ns | 22.9 | 2.1 | ns | 24.3 | 2.9 | ns | 4.5 | 1.1 | ns |
| 50nM ST | 0.9 | 0.4 | ns | 47.2 | 3.8 | ns | 23.0 | 2.2 | ns | 24.5 | 2.7 | ns | 4.4 | 0.8 | ns |
| 100nM ST | 2.3 | 0.8 | ** | 31.2 | 1.6 | **** | 26.7 | 2.1 | * | 35.8 | 3.9 | ** | 3.9 | 0.9 | ns |
| 300nM ST | 1.4 | 0.3 | ns | 20.6 | 1.3 | **** | 29.5 | 4.4 | ** | 44.8 | 5.1 | ** | 3.8 | 1.9 | ns |
| 10nM NOC | 1.2 | 0.2 | ns | 45.7 | 5.2 | ns | 22.1 | 2.1 | ns | 26.0 | 4.1 | ns | 5.0 | 0.8 | ns |
| 30nM NOC | 1.9 | 1.0 | * | 26.8 | 2.6 | ** | 27.3 | 2.4 | * | 37.5 | 6.5 | ** | 6.5 | 3.1 | ns |
| 50nM NOC | 1.6 | 0.9 | ns | 20.1 | 2.7 | ** | 29.0 | 4.5 | ** | 41.8 | 6.8 | ** | 7.5 | 2.8 | ** |
| 100nM NOC | 0.8 | 0.2 | ns | 13.8 | 3.4 | ** | 31.1 | 2.9 | ** | 47.4 | 1.3 | ** | 6.9 | 2.4 | * |
| 300nM NOC | 0.7 | 0.1 | ns | 14.5 | 2.7 | ** | 30.6 | 2.2 | ** | 46.6 | 1.1 | ** | 7.7 | 2.7 | ** |

**Supplementary Figure S3:** Cells exiting mitosis: NOC without cytokinesis and ST-401 with a lagging chromosome. Cell treated with CTR, NOC or ST (100 nM) were analyzed by flow cytometry using the classical FSC versus CCS signal to discriminate cell size. N = 3 independent experiments.

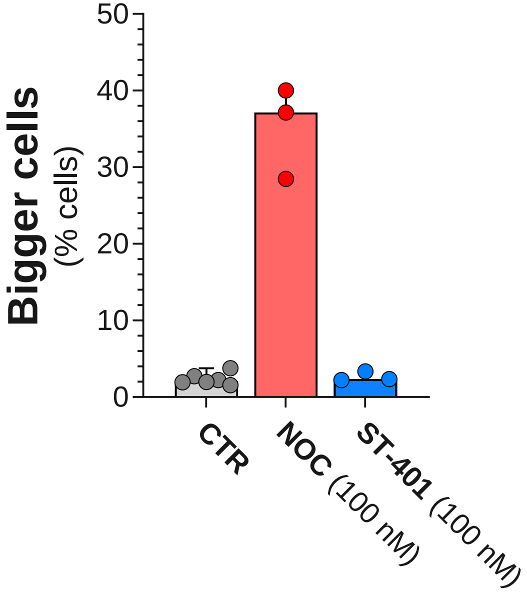

**Supplementary Figures S4a-c**: Quantification of MDC levels in HCT116 treated cells measured by flow cytometry. HCT116 cells treated with MTAs, harvested, and labeled with PI/Annexin V to measure apoptosis and necrosis using flow cytometry. Cells undergoing necrosis label A- and PI+, while labeling for A+ and PI- indicates early-apoptosis and A+/PI+ indicates late-apoptosis. Representative graphs of apoptosis/necrosis experiments at 100 nM. At 3 and 5 days we can see an increase in the number of cells at high FITC-A fluorescence, corresponding with high levels of Annexin V of cells going through apoptosis.

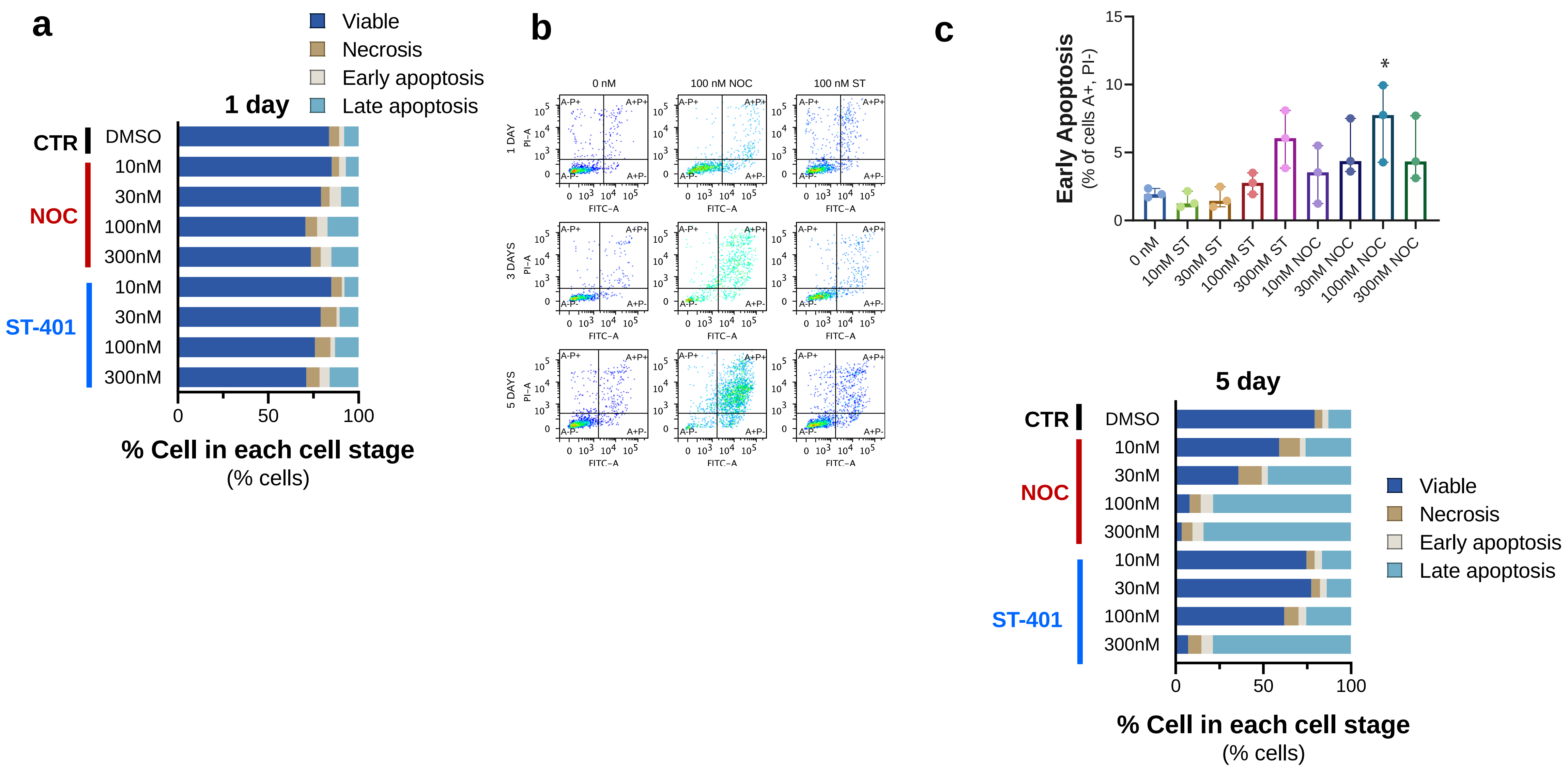

**Supplementary Figure S4d:** NOC trigger apoptosis: milder response by ST-401**.** Data were analyzed by ANOVA multi-comparison test followed by DUNETT's post test. N = 3 independent experiments. * = P < 0.0332, ** = P < 0.021, *** = P < 0.0002 and **** = P < 0.0001 different from Vehicle.

| 1 day | Viable | | | Necrosis | | | Early apoptosis | | | Late apoptosis | | |
|---|---|---|---|---|---|---|---|---|---|---|---|---|
| | Mean | SD | p | Mean | SD | p | Mean | SD | p | Mean | SD | p |
| DMSO | 83.62833 | 3.645125 | | 5.695 | 2.095898 | | 2.706667 | 1.533749 | | 7.973333 | 2.163235 | |
| 10nM ST | 84.92333 | 1.930423 | ns | 5.84 | 2.181582 | ns | 1.42 | 0.130767 | ns | 7.813333 | 0.550485 | ns |
| 30nM ST | 79.01 | 0.913947 | ns | 8.833333 | 2.306281 | ns | 1.533333 | 0.15308 | ns | 10.62 | 1.537173 | ns |
| 100nM ST | 75.78333 | 5.229812 | ns | 8.61 | 2.372109 | ns | 2.596667 | 0.67159 | ns | 13.01333 | 2.820591 | ns |
| 300nM ST | 71.00333 | 1.831784 | ** | 7.413333 | 1.59707 | ns | 5.496667 | 0.99143 | * | 16.08333 | 0.725695 | ** |
| 10nM NOC | 85.24667 | 4.737767 | ns | 3.943333 | 0.882968 | ns | 3.733333 | 1.916907 | ns | 7.076667 | 2.17238 | ns |
| 30nM NOC | 79.15667 | 4.259018 | ns | 4.833333 | 1.158462 | ns | 6.29 | 2.362478 | ** | 9.723333 | 1.348122 | ns |
| 100nM NOC | 70.57667 | 10.19993 | ** | 6.533333 | 1.767946 | ns | 5.693333 | 1.070389 | * | 17.19333 | 7.700424 | ** |
| 300nM NOC | 73.69667 | 1.260569 | * | 5.333333 | 1.568991 | ns | 5.903333 | 1.024809 | * | 15.06333 | 0.82282 | * |

| 3 days | Viable | | | Necrosis | | | Early apoptosis | | | Late apoptosis | | |
|---|---|---|---|---|---|---|---|---|---|---|---|---|
| | Mean | SD | p | Mean | SD | p | Mean | SD | p | Mean | SD | p |
| DMSO | 86.61 | 2.59513 | | 3.786667 | 1.155436 | | 1.996667 | 0.325167 | | 7.606667 | 1.669441 | |
| 10nM ST | 88.92667 | 3.2171 | ns | 3.886667 | 2.066454 | ns | 1.47 | 0.610492 | ns | 5.72 | 1.795188 | ns |
| 30nM ST | 87.06 | 2.781924 | ns | 3.97 | 1.51 | ns | 1.646667 | 0.754476 | ns | 7.33 | 2.536553 | ns |
| 100nM ST | 77.18333 | 6.473054 | ns | 6.226667 | 3.242628 | ns | 2.733333 | 0.785642 | ns | 13.85333 | 3.995552 | ns |
| 300nM ST | 22.33667 | 2.061173 | **** | 12.49 | 3.461344 | * | 6.003333 | 2.115569 | ns | 59.17667 | 4.139328 | **** |
| 10nM NOC | 61.51 | 9.880911 | ** | 12.62667 | 0.453468 | * | 3.43 | 2.137124 | ns | 22.43667 | 8.147443 | * |
| 30nM NOC | 40.32667 | 14.54225 | **** | 12.42667 | 5.645754 | * | 5.16 | 2.06182 | ns | 42.08 | 9.078128 | **** |
| 100nM NOC | 23.05333 | 4.275714 | **** | 10.73667 | 4.134941 | ns | 7.323333 | 2.854021 | * | 58.89667 | 5.965252 | **** |
| 300nM NOC | 11.26333 | 5.153565 | **** | 10.36333 | 3.14945 | ns | 5.053333 | 2.371863 | ns | 73.32333 | 5.237694 | **** |

| 5 days | Viable | | | Necrosis | | | Early apoptosis | | | Late apoptosis | | |
|---|---|---|---|---|---|---|---|---|---|---|---|---|
| | Mean | SD | p | Mean | SD | p | Mean | SD | p | Mean | SD | p |
| DMSO | 79.28333 | 1.696654 | | 4.403333 | 1.475545 | | 3.406667 | 1.301166 | | 12.90667 | 1.92962 | |
| 10nM ST | 74.54667 | 4.735149 | ns | 4.756667 | 1.030162 | ns | 4.013333 | 1.54442 | ns | 16.68667 | 2.804039 | ns |
| 30nM ST | 77.32667 | 2.916099 | ns | 4.996667 | 1.310432 | ns | 3.703333 | 1.720533 | ns | 13.97667 | 2.621475 | ns |
| 100nM ST | 61.97 | 9.258164 | ns | 8.093333 | 2.590103 | ns | 4.39 | 2.953032 | ns | 25.55 | 8.851107 | ns |
| 300nM ST | 7.09 | 0.818352 | **** | 7.613333 | 3.012812 | ns | 6.52 | 0.238118 | ns | 78.77333 | 2.906636 | **** |
| 10nM NOC | 59.07667 | 24.96963 | ns | 11.67667 | 8.382722 | ns | 3.353333 | 0.942992 | ns | 25.89333 | 16.83052 | ns |
| 30nM NOC | 35.80333 | 17.35193 | *** | 13.21333 | 4.825022 | ns | 3.5 | 0.915369 | ns | 47.48333 | 19.6458 | ** |
| 100nM NOC | 7.956667 | 0.647482 | **** | 6.406667 | 2.185002 | ns | 6.933333 | 1.254804 | ns | 78.70333 | 0.776938 | **** |
| 300nM NOC | 3.38 | 0.301993 | **** | 6.14 | 2.865118 | ns | 6.29 | 0.746458 | ns | 84.18667 | 2.917693 | **** |

**<u>Figure S5a-d</u>:** <u>Overview of single-cell RNA-seq sample metrics</u>. **a)** Boxplots of the number of unique molecular identifiers (UMIs) for the cells in our experiment. **b)** Violin plots of the percentage of mitochondrial reads per cell) for the cells in our experiment. **c)** Summary of the total cell number per sample. **d)** Correlation or replicate expression values (Pearson's rho).

**a**

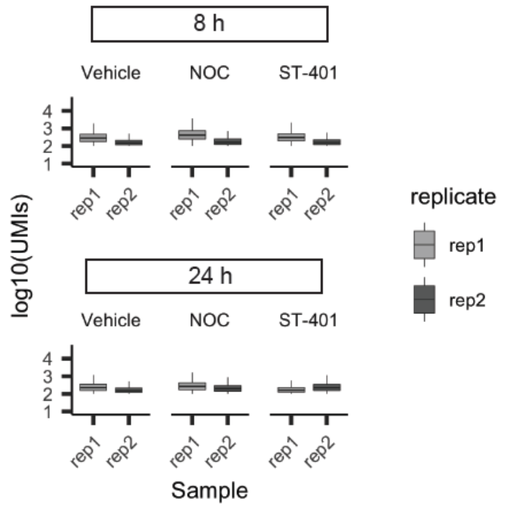


**b**

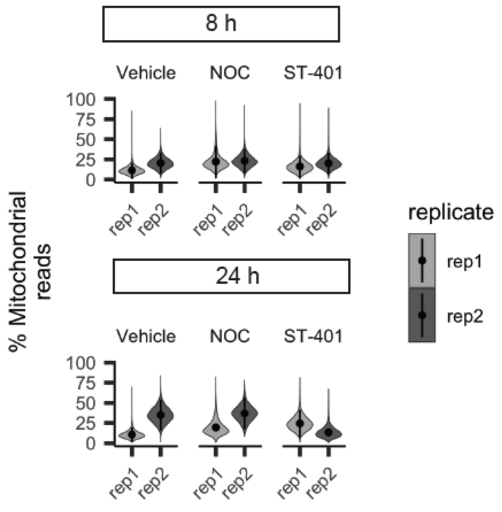


**c**

| Final cell counts (> 100 UMIs, < 25% mitochondrial reads) | | | |
|---|---|---|---|
| Treatment | Time point | Replicate | Total cells |
| Vehicle | 24 | Rep 1 | 43926 |
| Vehicle | 24 | Rep 2 | 2327 |
| Vehicle | 8 | Rep 1 | 29022 |
| Vehicle | 8 | Rep 2 | 6518 |
| NOC | 24 | Rep 1 | 33027 |
| NOC | 24 | Rep 2 | 219 |
| NOC | 8 | Rep 1 | 19124 |
| NOC | 8 | Rep 2 | 2768 |
| ST-401 | 24 | Rep 1 | 13431 |
| ST-401 | 24 | Rep 2 | 26854 |
| ST-401 | 8 | Rep 1 | 28907 |
| ST-401 | 8 | Rep 2 | 8042 |

**d**

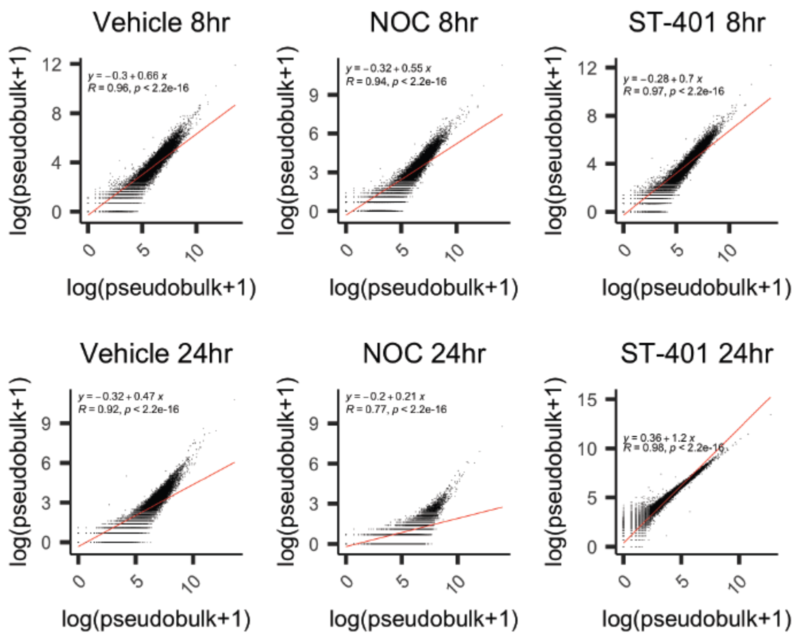

**Supplementary Figures S5e-f:** Summary of differential gene expression analysis of vehicle, NOC and ST-401 exposed cells. **e)** Volcano plots of the FDR and normalized $\beta_{coefficient}$ for the result of our differential gene expression analysis for every treatment and timepoint relative to vehicle control using a formula of "*expression* ~ *treatment* + *replicate*". The results of all test were combined and subjected to multiple hypothesis testing using the Benjamini-Hochberg procedure. **f)** UpSetR plot of the overlap of differentially expressed genes across each of our tests.

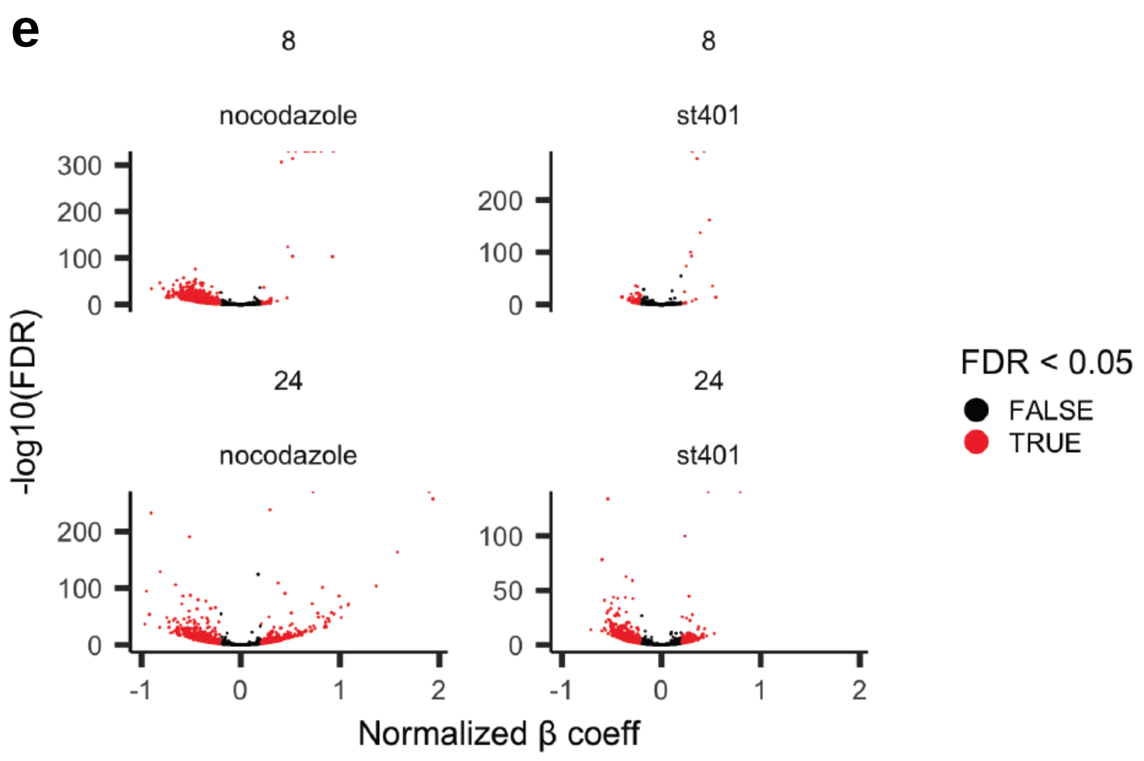


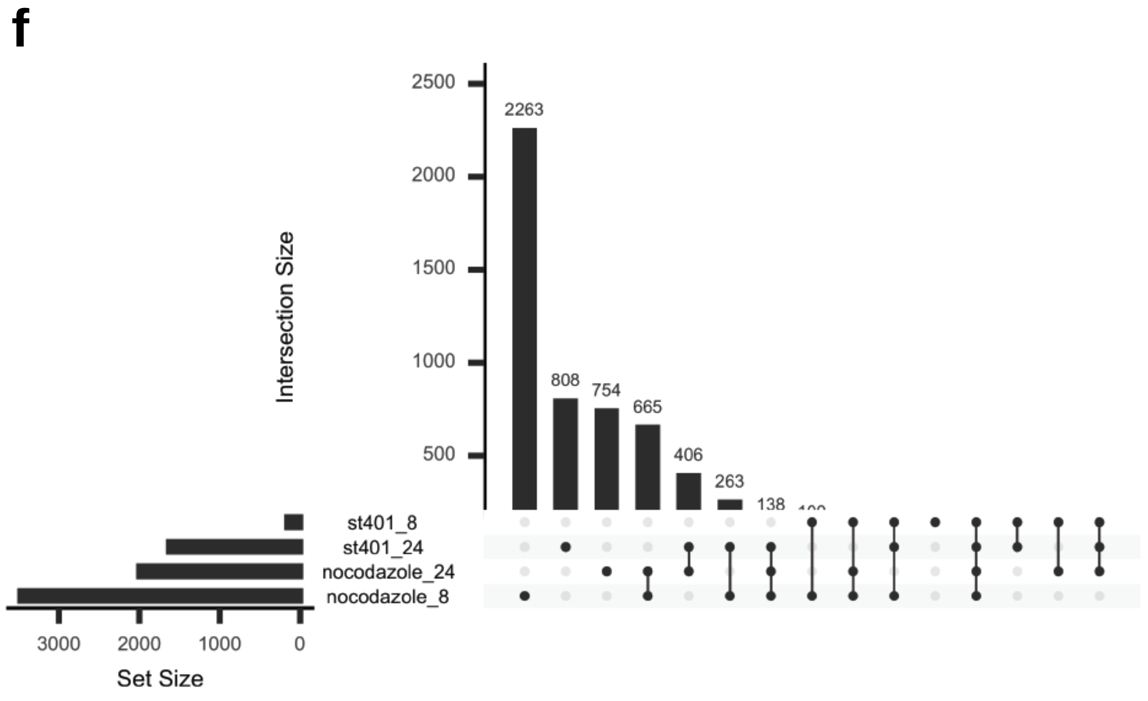

**Supplementary figures S5g-h:** Gene-enrichment analysis at 24h using Metascape. The different genes obtained from scRNAseq were divided in 4 quadrants depending on their expression levels under ST-401 and NOC treatment. The bioinformatics platform Metascape was used to obtain enrichment classifications per quadrant.

### $g_i$ Quadrant 1: ST-401↑ NOC↓

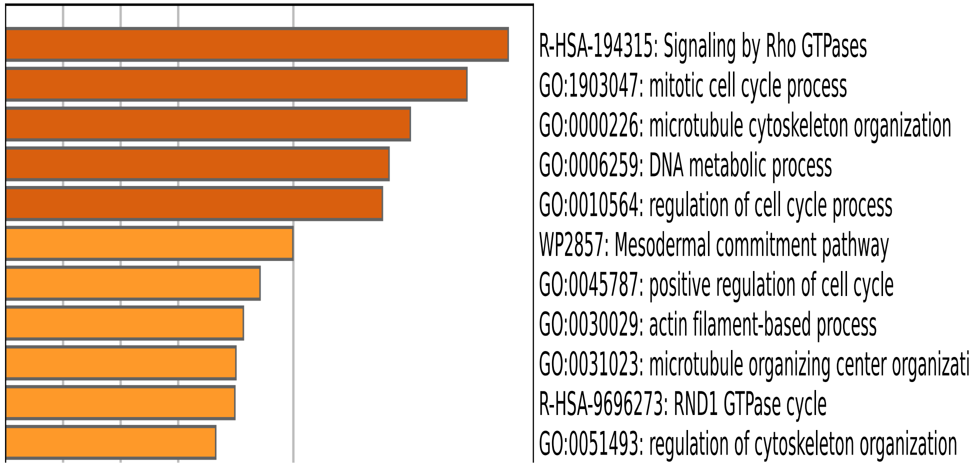


### $g_{ii}$ Quadrant 2: ST-401↑ NOC↑

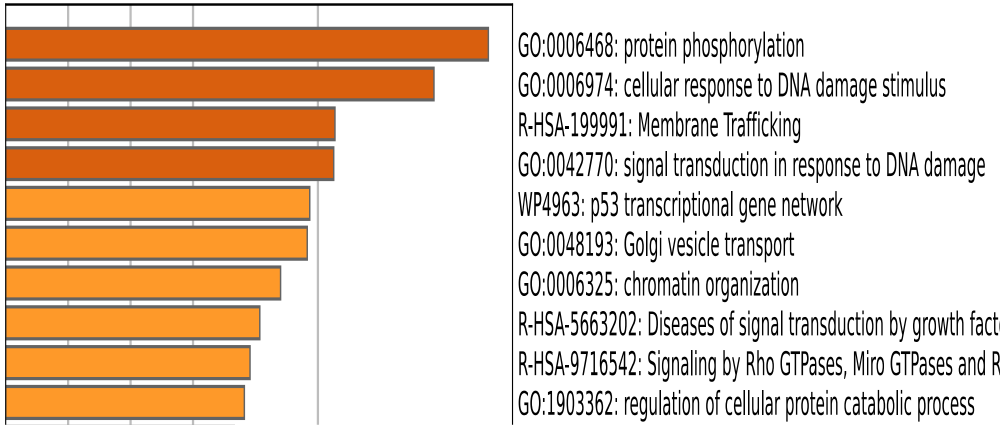


### $h_i$ Quadrant 3: ST-401↓ NOC↓

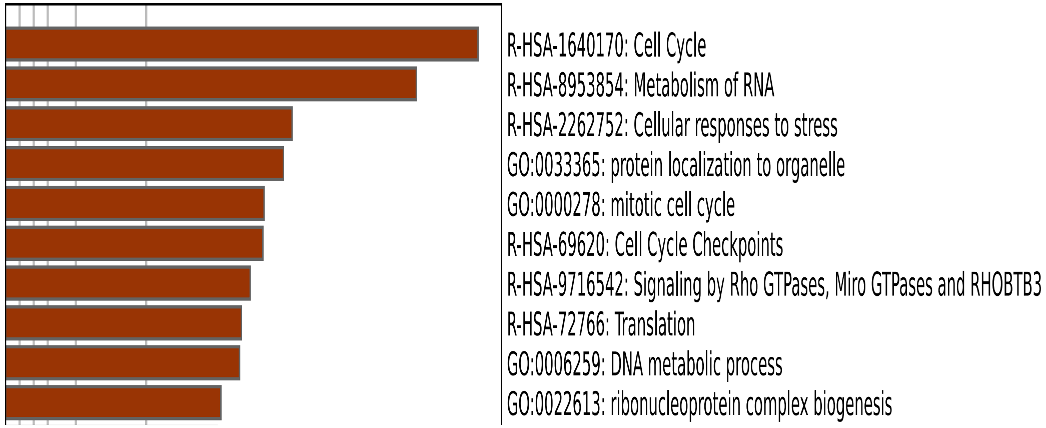


### $h_{ii}$ Quadrant 4: ST-401↓ NOC↑

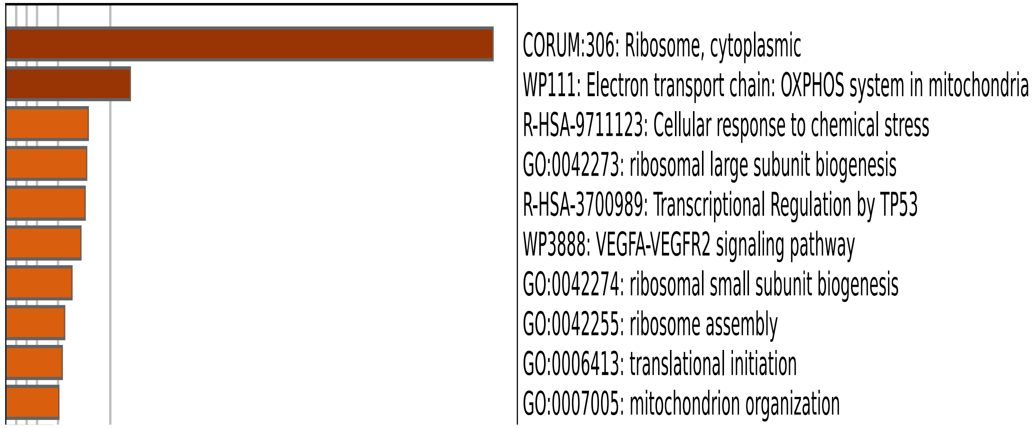

**Supplementary figure S5i:** Gene-enrichment analysis at 24h using Metascape. **i)** UMAP embedding of single-cell transcriptomes of cells exposed to DMSO (CTR), NOC, ST-401 for 8 (left) or 24 (right) h.

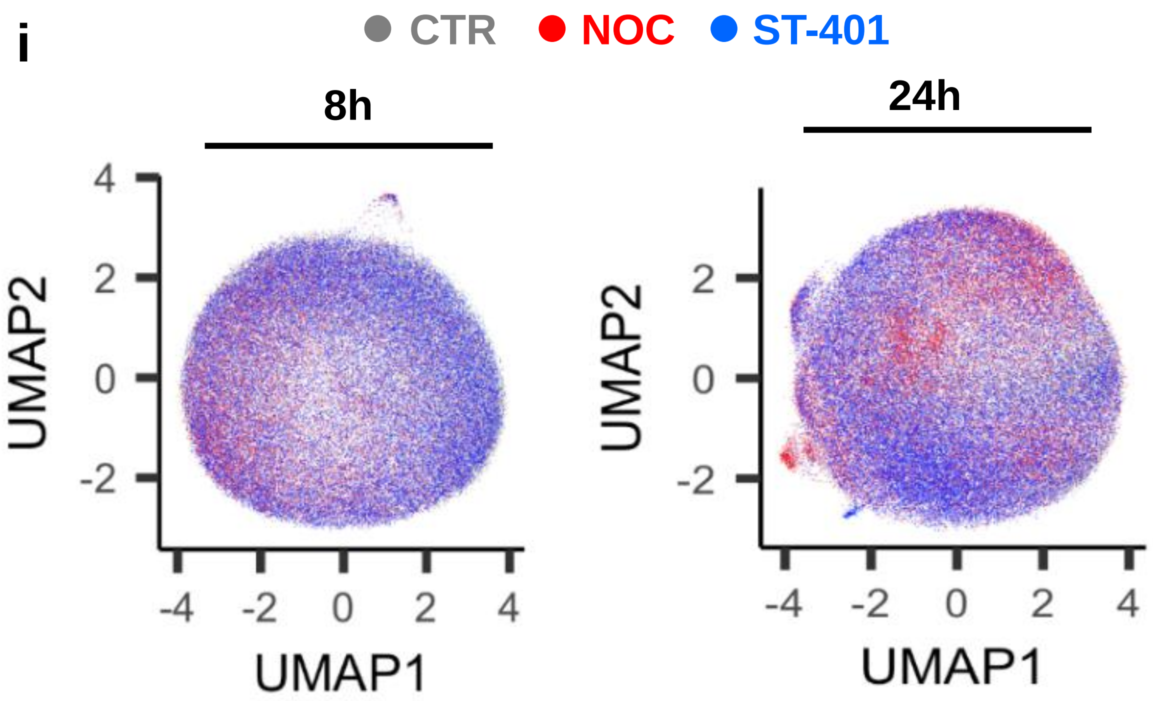

**Supplementary figure S6a-b:** Ponceau Red staining of gels shown in Figure 6. **a)** Ponceau Red stain of gel showing Puromycin analysis**. b)** Ponceau Red stain of gel showing Puromycin analysis.

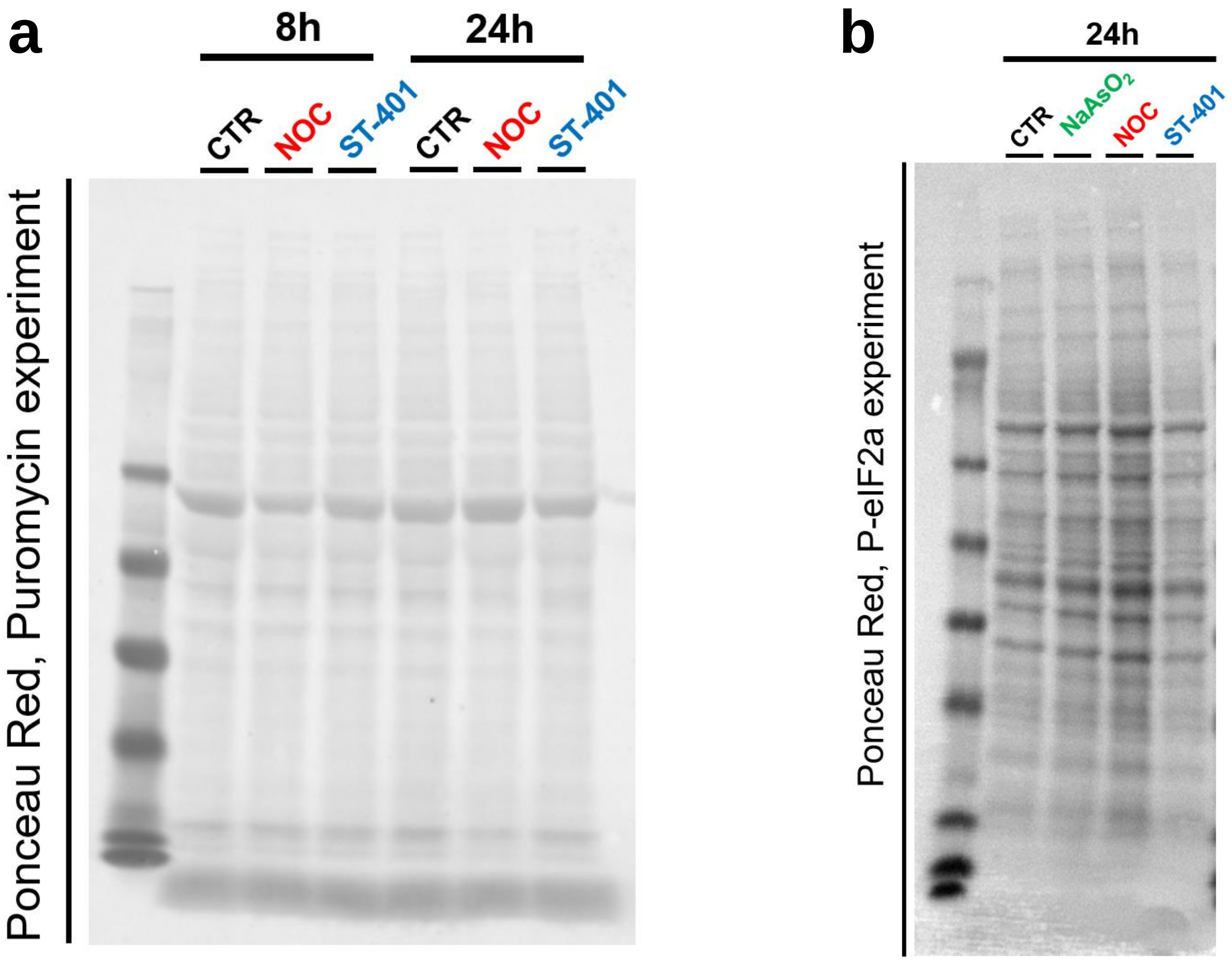

**Supplementary figure 7:** NOC (100 nM) and ST-401 (100 nM) treatment for 24h increases and decreases OCR and ECAR, respectively. The additional analyses that are linked to Figure 7. a) Additional OCR analyses and b) ECAR analyses.

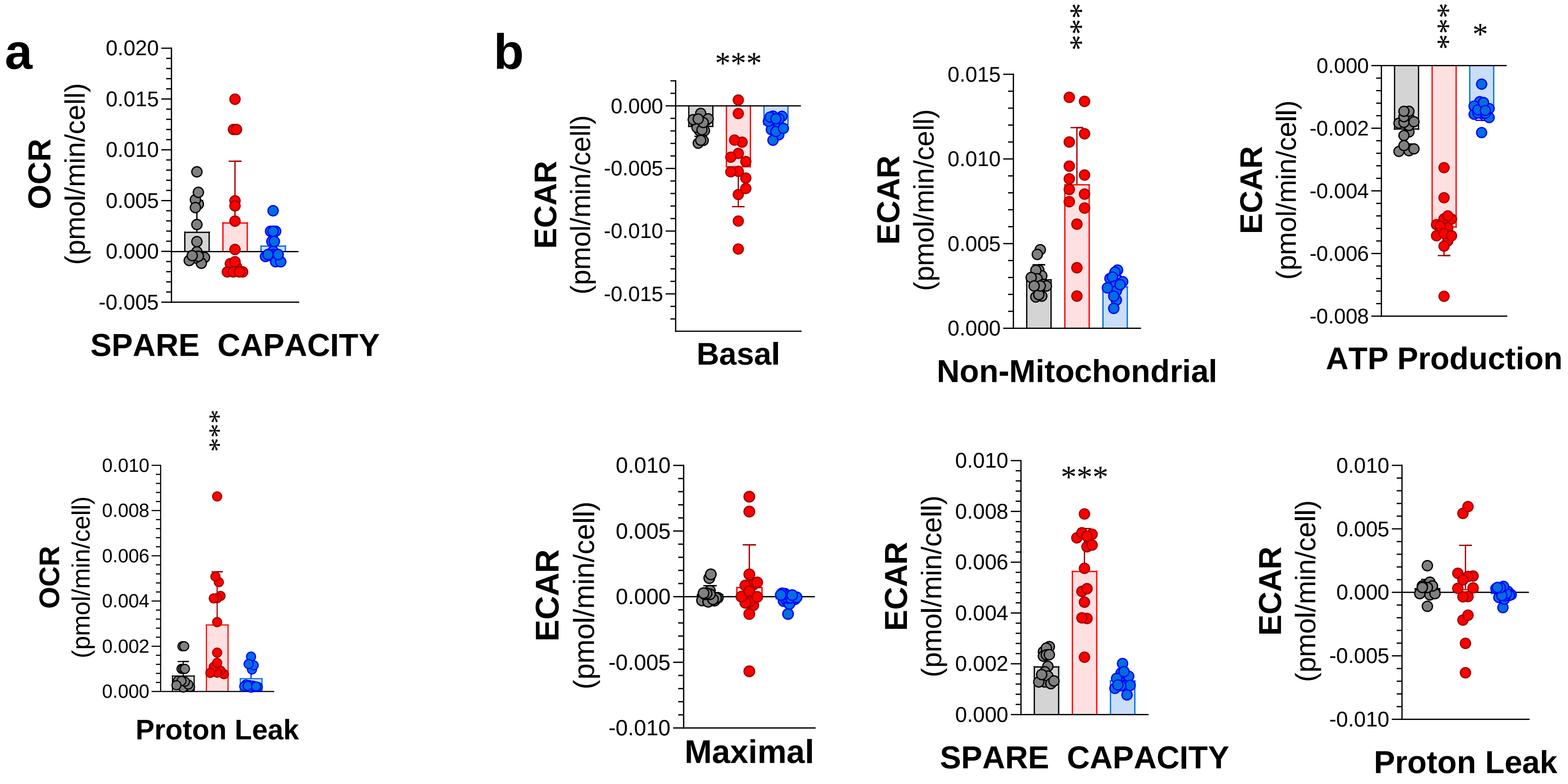

**Graphical Abstract:** Diagram depicting ST-401 antitumor activity. ST-401 treatment of HCT116 cells leads to death in interphase, transient reduction in protein synthesis, mitochondria fission and reduction in OXPHOS,

**2nd Graphical Abstracts that could be added, if asked by our editors and reviewers:** Diagram depicting that NOC treatment of HCT116 cells leads to death mitosis and apoptosis, autophagy, necrosis, sustained reduction in protein synthesis, mitochondria fusion and increase in OXPHOS. Cell that escape cell death become polyploid giant cancer cells.

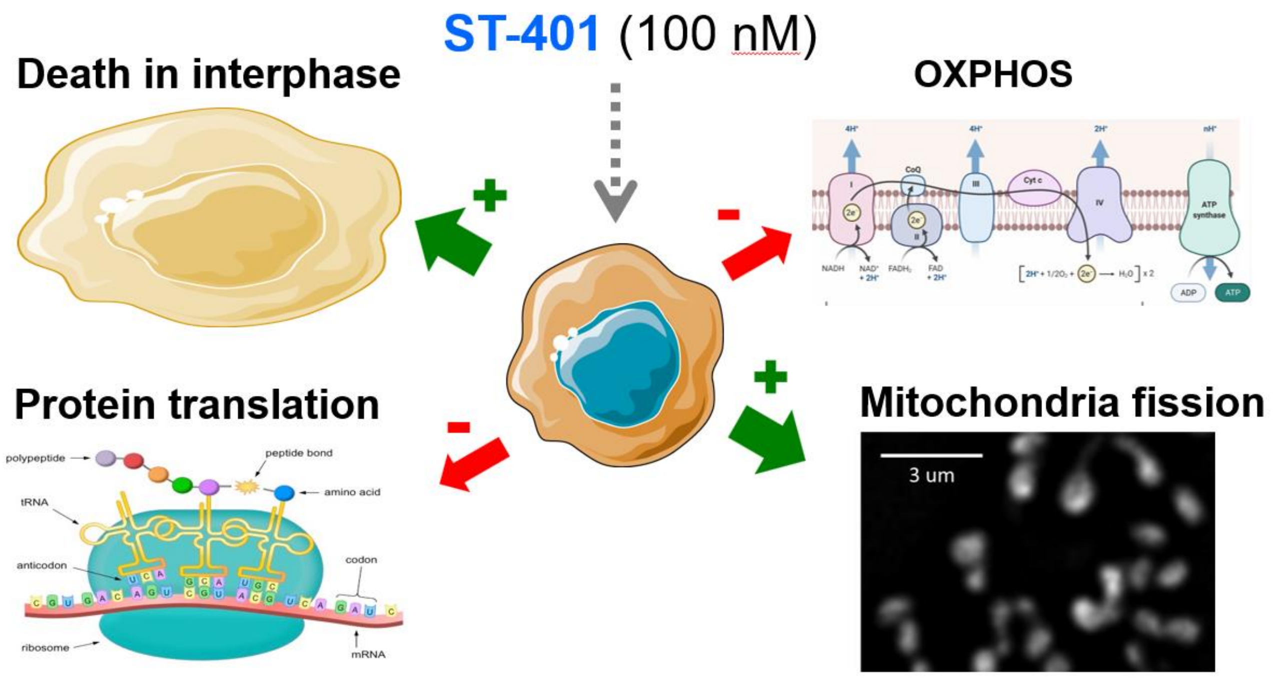


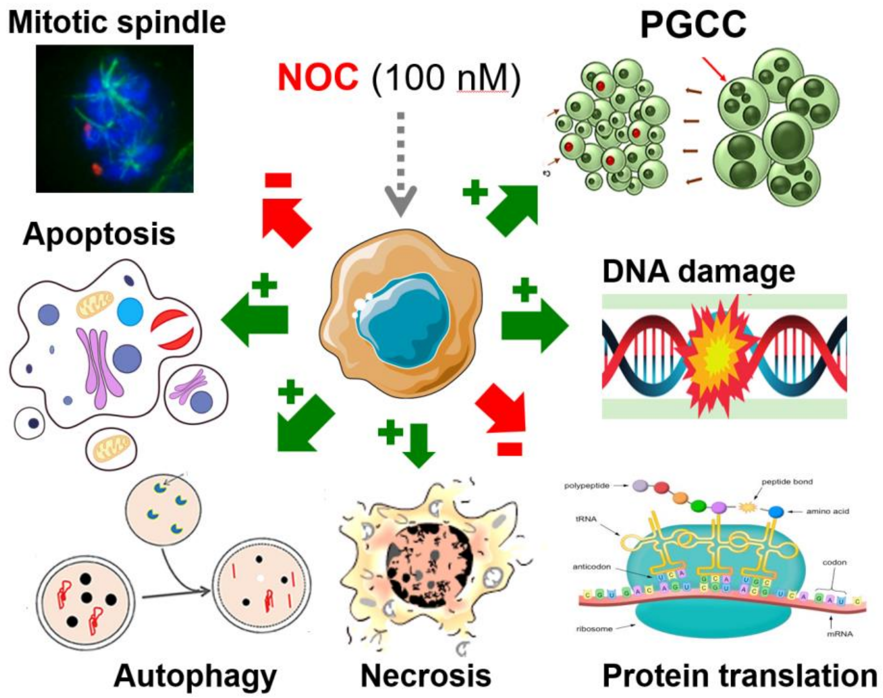
